# Inducible, virus-free direct lineage reprogramming enhances scalable generation of human inner ear hair cell-like cells

**DOI:** 10.1101/2025.02.20.639352

**Authors:** Robert N. Rainey, Sam D. Houman, Louise Menendez, Ryan Chang, Litao Tao, Helena Bugacov, Andrew P. McMahon, Radha Kalluri, John S. Oghalai, Andrew K. Groves, Neil Segil

## Abstract

Mammalian inner ear sensory hair cells are highly sensitive to environmental stress and do not regenerate, making hearing loss progressive and permanent. The paucity and extreme inaccessibility of these cells hinder the development of regenerative and otoprotective strategies. Direct lineage reprogramming to generate large quantities of hair cell-like cells in vitro offers a promising approach to overcome these experimental bottlenecks. Previously, we identified four transcription factors—*Six1*, *Atoh1*, *Pou4f3*, and *Gfi1* (SAPG)—capable of converting mouse embryonic fibroblasts, adult tail tip fibroblasts, and postnatal mouse supporting cells into induced hair cell-like cells through retroviral or lentiviral transduction (Menendez *et al*., 2020). Here, we developed a virus-free, inducible system using a stable human induced pluripotent stem (iPS) cell line carrying doxycycline-inducible SAPG. Our inducible system significantly increases reprogramming efficiency compared to retroviral methods, achieving a ∼19-fold greater conversion to a hair cell fate in half the time. Immunostaining, Western blot, and single-nucleus RNA-seq analyses confirm the expression of hair cell-specific markers and activation of hair cell gene networks in reprogrammed cells. The reprogrammed hair cells closely resemble developing fetal hair cells, as evidenced by comparison with a human fetal inner ear dataset. Electrophysiological analysis reveals that the induced hair cell-like cells exhibit diverse voltage-dependent ion currents, including robust, quick-activating, slowly inactivating currents characteristic of primary hair cells. This virus-free approach improves scalability, reproducibility, and the modeling of hair cell differentiation, offering significant potential for hair cell regenerative strategies and preclinical drug discovery targeting ototoxicity and otoprotection.

## Introduction

Hearing loss, primarily caused by the death of sensory hair cells in the inner ear, presents a global epidemic affecting hundreds of millions worldwide. Sensorineural hearing loss poses a significant challenge, with profound impacts on quality of life (Géléoc and Holt, 2014; Wong and Ryan, 2015). Unlike non-mammalian vertebrates, humans and other mammals lack the intrinsic ability to spontaneously regenerate lost hair cells in the cochlea, leading to progressive and irreversible deafness (Chardin and Romand, 1995). Many factors contribute to hearing loss, including genetic mutations, environmental stressors such as noise exposure and ototoxic drugs, as well as age-related degeneration (Matsui, Gale and Warchol, 2004; Wong and Ryan, 2015; Cheng, Cunningham and Rubel, 2005; Bodmer, 2008; Langer *et al*., 2013; Vaisbuch and Santa Maria, 2018). Current interventions for hearing loss primarily rely on prosthetic devices like cochlear implants and hearing aids, offering limited benefits for some individuals (Géléoc and Holt, 2014; Groves, 2010). Consequently, there is a pressing need to develop innovative strategies aimed at restoring or preserving cochlear structure and function, particularly through the regeneration of sensory hair cells or otoprotective drug screening. However, the complexities of the organ of Corti, composed of highly specialized hair cells and supporting cells, coupled with the paucity and inaccessibility of these cells, present substantial hurdles for regenerative medicine research and ototoxicity management (Chen and Segil, 1999; Lee, Liu and Segil, 2006; Ruben and Sidman, 1967; Lowenheim *et al*., 1999; Matei *et al*., 2005).

Efforts to address these challenges have led to the development of in vitro models capable of producing hair cell-like cells. Directed differentiation protocols, involving precise temporal manipulation of growth factors and signaling molecules within three-dimensional cultures, have enabled the generation of inner ear organoids from both embryonic stem cells and induced pluripotent stem cells (Koehler *et al*., 2013; Koehler *et al*., 2017; Moore *et al*., 2023; Jeong *et al*., 2018). However, the requirement for three-dimensional culture conditions and the variability in these approaches have constrained the feasibility of high-throughput studies with current imaging technologies. Alternatively, reprogramming with hair cell-specific transcription factors has emerged as a promising strategy for rapidly generating hair cell-like cells from readily available mammalian somatic cells (Iyer and Groves, 2021; Alonso *et al*., 2018; Iyer *et al*., 2022; Chen *et al*., 2021). In our previous study using direct lineage reprogramming, we successfully induced the conversion of mouse embryonic fibroblasts, adult tail tip fibroblasts, and postnatal mouse supporting cells into induced hair cells (iHCs) using *Six1*, *Atoh1*, *Pou4f3*, and *Gfi1* delivered by retroviral or lentiviral transduction (Menendez *et al*., 2020). We showed these hair cell-like cells to have similar transcriptional, epigenetic, morphological and electrophysiological properties to primary mouse hair cells. We further demonstrated that reprogrammed hair cells recapitulate the observed sensitivity of primary hair cells to gentamicin-induced ototoxicity. Recently, it was demonstrated that overexpression of *Atoh1*, *Pou4f3*, and *Gfi1* in nonsensory regions of the mature mouse cochlea induces the conversion of these cells into hair cell-like cells containing immature stereocilia bundles (McGovern *et al*., 2024). This suggests that although transcription factor reprogramming is feasible in the intact adult cochlea, further research is needed to understand the mechanism of reprogramming and address the obstacles in generating fully functional, mature hair cells.

While retroviral- and lentiviral-mediated reprogramming with hair cell-specific transcription factors is suitable for conducting high-throughput studies, its translational potential is constrained by several factors. The process of producing viruses, which constitutes the rate-limiting step, affects the scalability and robustness of robotic, high-throughput screening for drugs/small molecules capable of inducing hair cell regeneration, or protecting sensory hair cells from ototoxic agents. Infecting cells with multiple viruses is likely to strain cellular transcription/translation machinery (Babos *et al*., 2019), and only a subset of cells is expected to be infected by all the viruses (Mistry, D’Orsogna and Chou, 2018; Phan and Wodarz, 2015), thereby diminishing reprogramming efficiency. Additionally, certain cell types, such as induced pluripotent stem (iPS) cells, possess the ability to suppress viral expression after integration, significantly impeding the expression levels of the reprogramming factors.

To address these limitations, we have established a stable human iPS cell line carrying doxycycline-inducible *SIX1*, *ATOH1*, *POU4F3*, and *GFI1* reprogramming factors. We demonstrate the reliable and efficient reprogramming of this cell line to a hair cell-like state. Compared to retroviral-mediated methods, the virus-free induction system significantly enhances reprogramming efficiency and reproducibility, achieving a 19.1-fold increase in conversion efficiency in half the time. We show that reprogrammed hair cells closely resemble primary hair cells on the basis of their gene expression profiles and electrophysiological properties. The inducible system offers several advantages, including consistent expression of all four reprogramming factors, avoidance of viral silencing thus enabling the direct reprogramming of iPSCs and other stem cell types, and the ability to achieve transient expression of reprogramming factors, thereby better mimicking physiological hair cell differentiation. Leveraging human cells for this purpose also allows for modeling regenerative and otoprotective strategies unique to humans due to molecular differences between species. The increased efficiency and scalability of this approach provide a more robust tool for conducting high-throughput molecular and functional analyses. With access to increasingly larger gene and drug libraries, this advancement has the potential to significantly accelerate the screening process for genes that promote hair cell regeneration, as well as to facilitate pre-clinical discovery of drugs for combating ototoxicity and enhancing otoprotection.

## Results

### Reprogramming of human induced pluripotent stem (iPS) cells into hair cell-like cells (induced hair cells) using doxycycline-inducible *SIX1*, *ATOH1*, *POU4F3*, and *GFI1*

To improve upon the relatively low reprogramming efficiency inherent in transduction of multiple viruses in our previous strategy (Menendez *et al*., 2020), we developed a stable human iPS cell line carrying doxycycline-inducible hair cell reprogramming factors. This cell line harbors a doxycycline-inducible cassette expressing *SIX1*, *ATOH1*, *POU4F3*, and *GFI1* (collectively referred to as SAPG) from a single promoter. To create this cell line, we engineered a Tet-On inducible construct containing the SAPG reprogramming genes separated by 2A self-cleaving peptide sequences (Fig. 1*A*). The use of 2A self-cleaving peptides allows the generation of multiple proteins from a single primary transcript and ensures comparable expression levels of the reprogramming genes (Tang *et al*., 2009; Carey *et al*., 2009). We targeted the vector to the CLYBL safe harbor locus using CRISPR/Cas9-mediated knock-in (Lo, Greben and Chen, 2017) (Fig. 1*A*). Additionally, the cell line carries a *POU4F3*-tdTomato hair cell reporter integrated independently at a separate locus in the genome, enabling live tracking of reprogrammed hair cells. This reporter is not expressed in the starting iPSC population and becomes activated as hair cell-like cells begin to differentiate, approximately 3 days after doxycycline administration (Fig. 1*B*; Supp. Fig. 1).

**Figure 1.**
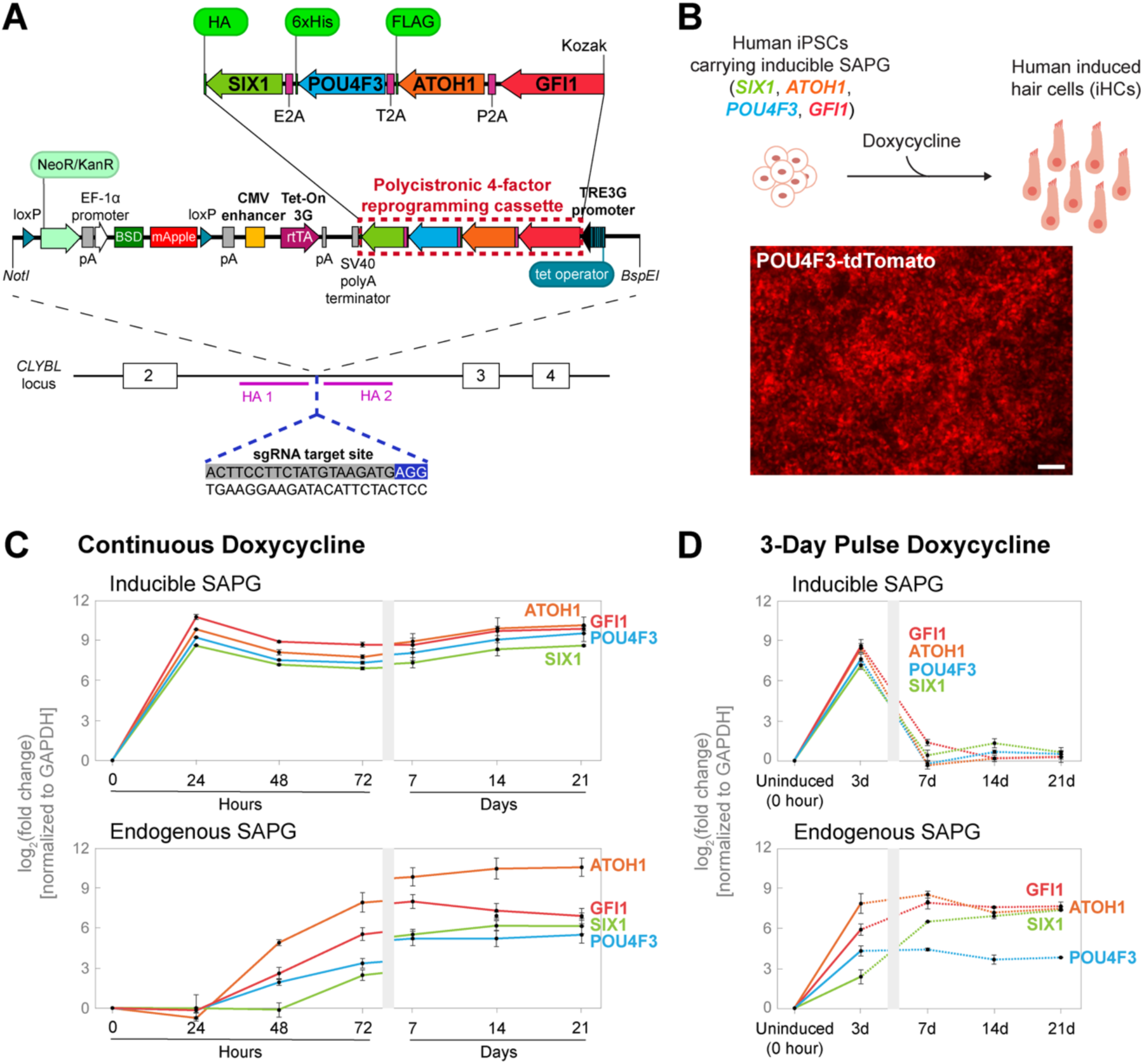
The stable human induced pluripotent stem (iPS) cell line carrying doxycycline-inducible *SIX1*, *ATOH1*, *POU4F3*, and *GFI1* promotes robust induction of hair cell transcription factors. (*A*) Schematic representation of the strategy used to generate the stable human iPS cell line capable of inducible expression of hair cell transcription factors (TFs). A Tet-On vector containing polycistronic four-TF (*SIX1*, *ATOH1*, *POU4F3*, and *GFI1*) reprogramming cassette (highlighted by red dashed box) under the control of a TRE3G inducible promoter is shown at the top. The four TFs were separated by 2A self-cleaving peptides to generate multiple proteins from a single primary transcript. Epitope tags were incorporated at the 3’ end as follows: the HA tag with *SIX1*, the 6xHis tag with *POU4F3*, and the FLAG tag with *ATOH1*. The vector was integrated into the CLYBL safe harbor locus between the two homology arms (HA1 and HA2) using CRISPR/Cas9-mediated knock-in. Upon confirmation of correct targeting, the cell line was subsequently treated with TAT-CRE recombinase to excise the mApple screenable marker, FACS-sorted as mApple-negative cells, followed by clonal expansion. (*B*) Schematic of direct reprogramming of human iPSCs into hair cell-like cells (induced hair cells) using the doxycycline-inducible system. The *POU4F3*-tdTomato hair cell reporter becomes visible after ∼3 days of reprogramming. Representative image shown was acquired at 7 days post-doxycycline treatment. Scale bar represents 200 µm. (*C*-*D*) RT-qPCR analysis of the cell line treated with continuous doxycycline (*C*) or a 3-day pulse of doxycycline (*D*) over the indicated time points. Dashed lines indicate post-doxycycline removal. Values are normalized to GAPDH, and a ratio is calculated by dividing the uninduced control (0 hour). Fold-change values are log2-transformed. Error bars indicate SEM. *n* = 3 biological replicates. Statistical analyses are provided in Supplementary file 1.

We tested the ability of the stable cell line to express individual reprogramming genes (Inducible SAPG) and their endogenous counterparts (Endogenous SAPG) over time in the presence of continuous doxycycline treatment (Fig. 1*C*). Using RT-qPCR and primer sets designed to distinguish between SAPG transgenes and endogenous genes, we observed robust induction (log_2_[fold change] of 8.58–10.78) of the reprogramming factors 24 hours after doxycycline administration. Expression of the reprogramming factors remained consistently elevated for up to 21 days, the longest time point tested (Fig. 1*C*). In contrast, endogenous SAPG genes were expressed around ∼48 hours after doxycycline administration and reached steady-state levels by 7 days, persisting through to 21 days. We also investigated the responsiveness of the inducible system to either a 3-day (Fig. 1*D*) or a 7-day (Supp. Fig. 2) pulse of doxycycline.

Following the 3- or 7-day pulse, we observed significant induction of the reprogramming factors, with their expression returning to near or at baseline levels by day 7 (after 3-day pulse; Fig. 1*D*) or by day 14 (after 7-day pulse; Supp. Fig. 2). Notably, pulsed doxycycline administration activated endogenous *GFI1* and *SIX1* expression to levels comparable to those observed under chronic doxycycline condition. In contrast, endogenous *ATOH1* and *POU4F3* expression showed a decrease but still maintained strong induction by day 21, with log_2_[fold change] values of 7.49 ± 0.15 and 3.83 ± 0.03, respectively (after 3-day pulse; Fig. 1*D*) or 8.28 ± 0.14 and 4.44 ± 0.23, respectively (after 7-day pulse; Supp. Fig. 2).

To assess the efficacy of the doxycycline-inducible system in reprogramming human iPSCs to a hair cell-like state, we performed immunostaining for established hair cell-specific markers (POU4F3, MYO7A, and MYO6; Fig. 2 *A* and *B*; Supp. Fig. 3) and the pluripotency marker NANOG (Fig. 2*A*). We observed staining for POU4F3 and MYO7A, and activation of the *POU4F3*-tdTomato reporter (Supp. Fig. 1), three days after initiating doxycycline treatment, coinciding with the loss of iPSC identity, as indicated by the disappearance of NANOG (Fig. 2*A*). After seven days of continuous doxycycline, 58.6% (± 0.3%) of cells were positive for transgene encoded, HA epitope-tagged SIX1, while 34.1–40.1% were reprogrammed to a hair cell-like state, as quantified by the presence of POU4F3+, MYO7A+, or MYO6+ cells (Fig. 2*B*; Supp. Fig. 3). In contrast, only 2.1% (± 0.2%) of cells tested positive for MYO7A when reprogrammed via retroviral-mediated transduction over 14 days, the standard duration used in our previous study (Menendez *et al*., 2020) (Fig. 2*C*). This reflects a 19.1-fold increase in the conversion efficiency to MYO7A+ cells, achieved by the inducible system in half the time required for retroviral-mediated reprogramming (Fig. 2*C*). RT-qPCR analysis revealed that *MYO7A* and *MYO6* were activated by day 3 and reached peak expression starting after 7 days of reprogramming in continuous doxycycline (Fig. 2*F*). Western blot analysis confirmed the robust induction and maintenance (day 14–21) of MYO7A and MYO6 and loss of pluripotency associated POU5F1 (OCT4) in the presence of doxycycline (Fig. 2 *D* and *E*). Importantly, *MYO7A* and *MYO6* expression was sustained after withdrawal of doxycycline at day 3 (Fig. 2*G*) or day 7 (Supp. Fig. 2), albeit at reduced levels, consistent with transcriptional regulation by endogenous *ATOH1* and *POU4F3* (Fig. 1*D*; Supp. Fig. 2).

**Figure 2.**
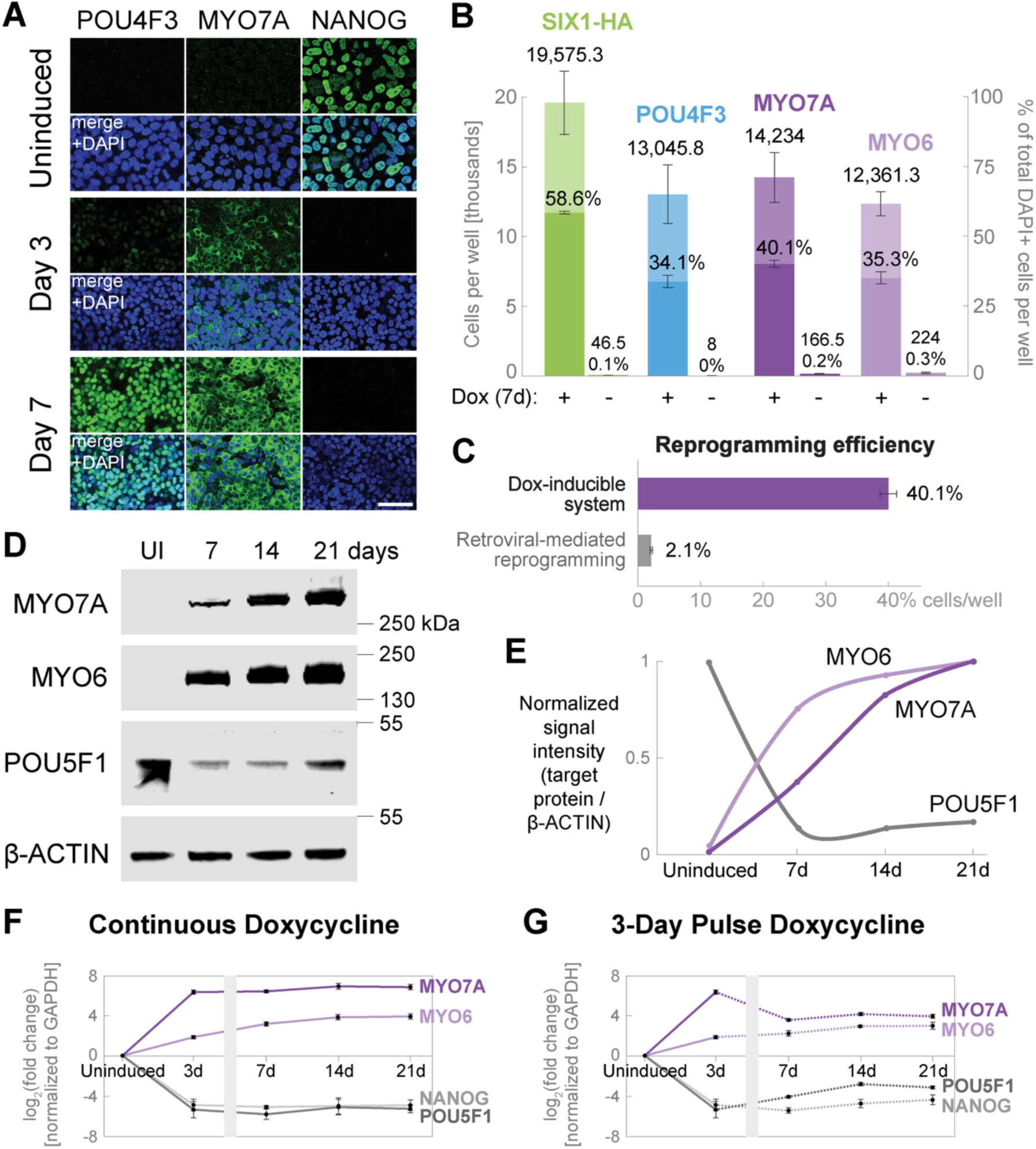
The stable cell line enables highly efficient reprogramming of human iPSCs into hair cell-like cells via inducible expression of *SIX1*, *ATOH1*, *POU4F3*, and *GFI1*. (*A*) Representative images of uninduced control and after 3 or 7 days of reprogramming with continuous doxycycline treatment. Immunostaining for hair cell markers POU4F3 and MYO7A coincides with the loss of pluripotency marker NANOG, indicating the conversion of iPSCs into hair cell-like cells. Scale bar represents 50 µm. (*B*) Quantification of human induced hair cells after 7 days of reprogramming. Cells were plated in a 96-well format and treated either with continuous doxycycline for 7 days (+Dox) or left untreated (−Dox) and doubled-labeled with DAPI nuclear stain and antibodies against the HA epitope tag, POU4F3, MYO7A, or MYO6. The HA tag, attached to the 3’ end of the *SIX1* transgene in the reprogramming cassette (see Fig. 1A), serves as a representative marker to track the expression of reprogramming factors. The presence of each marker was quantified from *n* = 4 wells, represented as total number of cells per well (left y-axis) and as a percentage out of the total number of DAPI+ cells per well (right y-axis). Data are presented as mean ± SEM. (*C*) Comparison of conversion efficiency between the doxycycline-inducible system (7 days of reprogramming) and retroviral-mediated transduction (14 days of reprogramming, the standard duration used in our previous study (Menendez *et al*., 2020)). Compared to retroviral reprogramming, the inducible system exhibits a 19.1-fold increase in conversion efficiency to MYO7A+ cells per well of a 96-well plate. Error bars indicate SEM. Reprogramming efficiencies were compared using an unpaired two-tailed Welch’s *t*-test. *p* < .0001. (*D*-*E*) Western blot analysis of uninduced control (UI) and cells treated with continuous doxycycline for 7, 14, or 21 days. Protein abundance was quantified by normalizing the signal intensity of each target protein to β-ACTIN and graphed on a scale of 0 to 1 along the y-axis. (*F*-*G*) RT-qPCR analysis of cells treated with continuous doxycycline (*F*) or a 3-day pulse of doxycycline (*G*) over the indicated time points. Dashed lines indicate post-doxycycline removal. Values are normalized to GAPDH, and a ratio is calculated by dividing the uninduced control (0 hour). Fold-change values are log_2_- transformed. Error bars indicate SEM. *n* = 3 biological replicates. Statistical analyses for (*B*-*C*) and (*F*-*G*) are provided in Supplementary file 1.

### Single-nucleus RNA-seq profiling reveals hair cell gene networks

To gain further insight into the direct reprogramming of human iPSCs into hair cell-like cells, we treated cells with doxycycline continuously for 21 days then performed single-nucleus RNA sequencing (snRNA-seq) and “uniform manifold approximation and projection” (UMAP) to compare with uninduced cells (Fig. 3*A*). Uninduced (6,048 nuclei) and induced (5,028 nuclei) samples segregated into three and five clusters, respectively (Fig. 3*B*). As expected, the major uninduced cluster (labeled Control cluster 1 [C1]) showed typical features of pluripotency such as *POU5F1* expression and cell cycle progression (*CCND1*, *CCNB1*, *CDK1*, *BIRC5)* (Fig. 3*B*, *D*-*E*). Among the five clusters from day 21 reprogrammed samples, Reprogrammed cluster 5 (R5) expressed established hair cell markers (*MYO7A*, *MYO6*, *MYO15A*, *CCER2*, *KCNH6*, *LHX3*) at significantly higher levels and in a larger percentage of cells compared to other clusters (R1–4; Fig. 3*B*, *D*-*E*). This cluster also exhibited robust activation of both inducible and endogenous SAPG, with higher endogenous expression levels of *ATOH1* and *POU4F3* compared to *SIX1* and *GFI1* (Fig. 3*C*).

**Figure 3.**
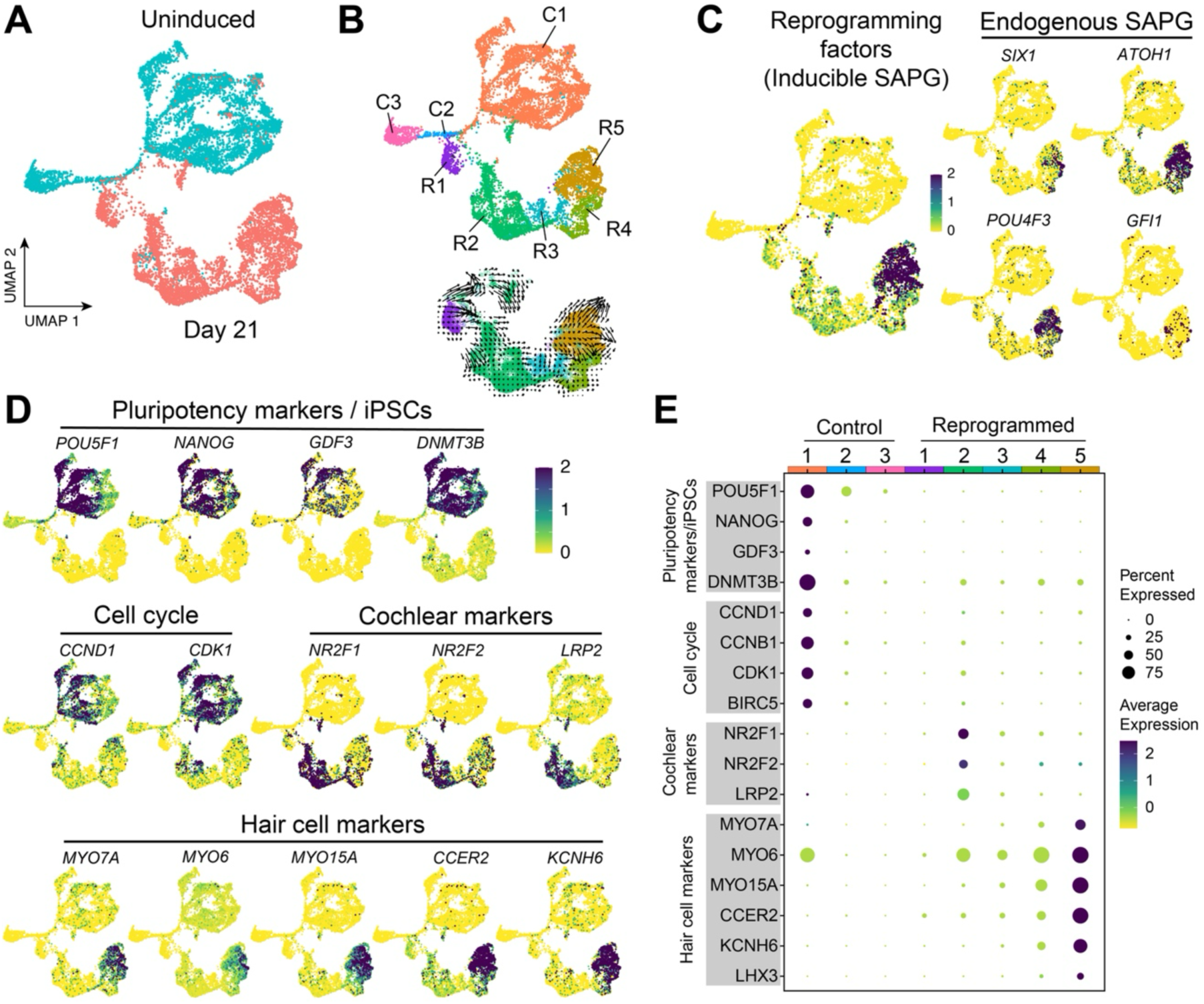
Single-nucleus transcriptional profiling of the stable cell line demonstrates hair cell gene network activation following inducible expression of *SIX1*, *ATOH1*, *POU4F3*, and *GFI1*. (*A*) Uninduced control and cells treated continuously with doxycycline for 21 days were collected and separately profiled using single-nucleus RNA sequencing (snRNA-seq). Integration of these datasets and subsequent UMAP projection after unbiased Seurat clustering revealed a clear separation between the two conditions (Uninduced and Day 21). (*B*) Clustering analysis identified three distinct clusters for uninduced control (C1–3), while five distinct clusters were observed for reprogrammed cells (R1–5). RNA velocity (trajectory) of reprogrammed cells (clusters R1–5) was visualized using Velocyto.R. (*C*) Among the reprogrammed clusters, higher activation levels of SAPG (both inducible SAPG and endogenous SAPG) were observed in cluster R5 compared to the other four reprogrammed clusters (R1–4). (*D*-*E*) Expression of markers associated with pluripotency (*POU5F1*, *NANOG*, and *GDF3*), or the iPSC state (*DNMT3B*) (Chan *et al*., 2009), and genes involved in cell cycle progression, was predominantly found in cluster C1. Conversely, cochlear marker genes (*NR2F1/2* and *LRP2*) were activated at higher levels in cluster R2, while hair cell markers were expressed at higher levels in cluster R5. The gene expression dataset associated with this figure is available in <u>gEAR</u>.

We interrogated the reprogrammed populations for other genes associated with inner ear cell identity. Interestingly, the cochlear markers *NR2F1/2* and *LRP2* were upregulated in cluster R2 but expressed at lower levels in the remaining clusters (Fig. 3 *D* and *E*). *NR2F1/2* are highly differentially expressed in cochlear but not vestibular organoids (Moore *et al*., 2023). Likewise, *NR2F1* is expressed in cochlear hair cells but absent in vestibular hair cells in human fetal cochleae at gestation week 18 (Moore *et al*., 2023). Similarly, *LRP2*, a downstream effector in the Sonic Hedgehog (SHH) pathway necessary for cochlear induction, is enriched in ventralized otic progenitors (Moore *et al*., 2023) as well as in the marginal cells of the stria vascularis (Faridi *et al*., 2023). We did not detect significant activation of markers associated with supporting cell identity in any of the reprogrammed cell clusters (Supp. Fig. 4). This suggests that reprogramming with SAPG does not activate supporting cell networks in the stable cell line.

### Retroviral-mediated transduction of human fibroblasts with *SIX1*, *ATOH1*, *POU4F3*, and *GFI1* demonstrates comparable reprogramming fidelity

Given the significantly higher reprogramming efficiency observed in the stable cell line compared to retroviral-mediated reprogramming with SAPG (Fig. 2*C*), we wondered whether the doxycycline-inducible system also enhances the fidelity of hair cell reprogramming by activating a greater number of hair cell genes. To investigate this, we performed retroviral-mediated reprogramming of human iPSC-derived fibroblasts with SAPG over a 21-day period. To do this, we utilized the parent line (carrying the *POU4F3*-tdTomato reporter but lacking the reprogramming cassette) of the stable cell line. Before initiating reprogramming, we converted the iPSCs into secondary fibroblasts, a necessary step due to retroviral silencing in iPSCs. We profiled sorted populations of *POU4F3*-tdTomato^+^ and *POU4F3*-tdTomato^−^ cells (in a single sample) collected after 21 days of reprogramming using single-cell RNA-seq. Of the 6,676 cells analyzed, we found reprogramming hair cell-like cells to be segregated into three distinct clusters, as depicted by UMAP projection (Fig. 4*A*). These clusters coincided with canonical hair cell markers, including *MYO6* and *MYO15A* (Fig. 4 *A* and *B*; Supp. Fig. 5), leading us to label them Retrovirus-Reprogrammed clusters 1 through 3 (RV-R1–3; Fig. 4*A*). In contrast, the expression of *CCND1*, which encodes cyclin D1 involved in cell cycle progression, and the mesenchymal marker *COL2A1*, was evident in residual fibroblasts (Fig. 4*A*). Additionally, the reprogrammed clusters exhibited reduced expression of *NOTCH1/2/3*, and upregulation of hair cell-specific NOTCH ligand genes *DLL3*, *JAG2*, and *DLK2*, relative to fibroblast clusters (Supp. Fig. 6). This is consistent with the reported modulation of NOTCH signaling pathway during hair cell differentiation and maturation (Kelley, 2006). The observed trajectory in the RNA velocity map (Fig. 4*A*), and heat map (Fig. 4*B*; Supp. Fig. 5), indicate that clusters 2 and 3 are more hair cell-like compared to cluster 1. Pseudotime analysis of clusters 1–3 using Monocle (Qiu *et al*., 2017) demonstrates that reprogramming cells in clusters 1 and 2 are still in the process of erasing fibroblast identity, characterized by the downregulation of fibroblast-enriched genes *GYPC*, *CPXM1*, *COL5A2*, and *SDC2* (Supp. Fig. 7). In contrast, clusters 2 and 3 adopt a hair cell gene expression signature, exemplified by the upregulation of *TNC*, *MREG*, *COL9A2*, and *DLK2* (Supp. Fig. 7).

**Figure 4.**
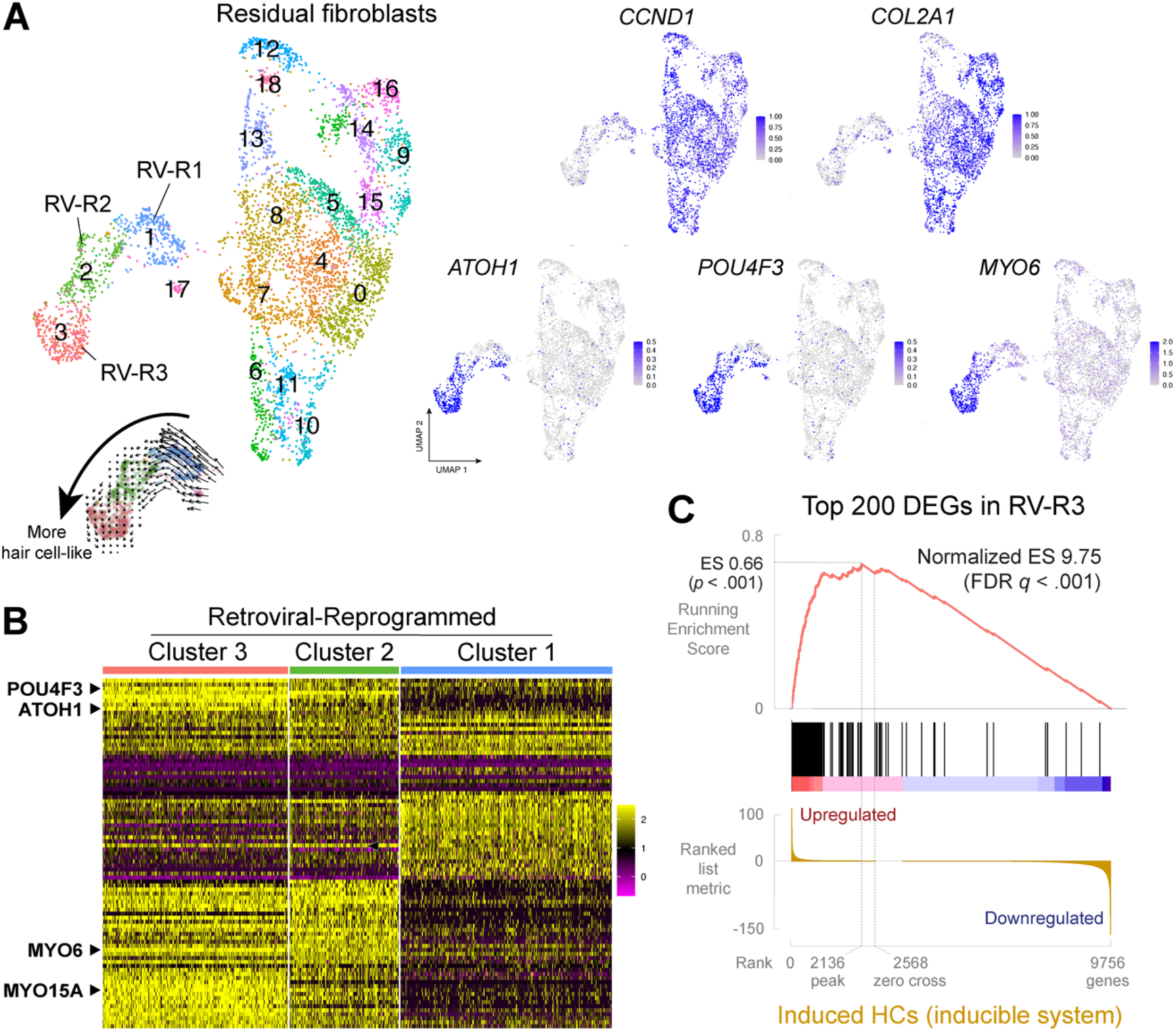
*SIX1*, *ATOH1*, *POU4F3*, and *GFI1* effectively reprogram human fibroblasts to a hair cell-like state via retroviral transduction, exhibiting activation of hair cell gene networks comparable to that observed with the inducible system. (*A)* Human iPSC-derived fibroblasts were reprogrammed with a cocktail of retroviruses expressing *SIX1*, *ATOH1*, *POU4F3,* or *GFI1* as described in (Menendez *et al*., 2020) for 21 days. A sorted population consisting of 50% *POU4F3*-tdTomato^+^ cells and 50% *POU4F3*-tdTomato^−^ cells was profiled using single-cell RNA- seq and plotted using UMAP projection. RNA velocity of hair cell-like clusters (labeled Retrovirus- Reprogrammed clusters 1 through 3 [RV-R1–3]) was visualized using Velocyto.R, indicating a transition of reprogramming cells from cluster 1 to cluster 3. (*B*) Heat map of the top 30 differentially expressed (DE) genes present in each of the hair-cell like clusters (RV-R1–3). (*C*) Gene set enrichment analysis (GSEA) was performed on the top 200 DE gene set identified in RV-R3 against cluster R5 (Fig. 3) generated by the doxycycline-inducible system. 86.7% of the gene set identified in SAPG-transduced hair cell-like cells was also activated in hair cell-like cells reprogrammed from the stable cell line. The ranked list metric is sorted by log_2_(fold change)* −log_10_(adj *p*-value) (shown in thousands). The gene expression dataset associated with this figure is available in <u>gEAR</u>.

Using gene set enrichment analysis (GSEA) (Subramanian *et al*., 2005), we examined whether the top differentially expressed (DE) genes in the most hair cell-like cluster 3 (RV-R3) were enriched in reprogrammed hair cells generated by the inducible system (cluster R5). The top 200 DE gene set identified in RV-R3 was substantially enriched in R5, with a normalized enrichment score of 9.75 at FDR (false discovery rate) *q* < 0.1% (Fig. 4*C*). This finding indicates that 86.7% of the gene set identified in SAPG-transduced hair cell-like cells was also activated in hair cell-like cells reprogrammed from the stable cell line. Additionally, reprogrammed hair cells derived from both human and mouse fibroblasts exhibit significant enrichment of each other’s top DE genes, with a normalized enrichment score of 8.20 (human iHCs) or 7.31 (mouse iHCs) (both FDR *q* < 0.1%; Supp. Fig. 8). These results suggest that, while the doxycycline-inducible system enhances the efficiency of hair cell reprogramming, it does not notably improve the fidelity of reprogramming compared to retroviral-mediated transduction with SAPG.

### Reprogramming with *SIX1*, *ATOH1*, *POU4F3*, and *GFI1* produces human iHCs with a transcriptome similar to fetal human hair cells and mixed hair cell subtype expression patterns

We focused on the successfully reprogrammed hair cell-like cells for further characterization. To offer a reference of human primary hair cells for comparative analysis, we subsetted C1 and R5 (representing iPSCs and reprogrammed hair cells, respectively; Fig. 3) from our dataset and integrated them with the human fetal inner ear snRNA-seq dataset published in van der Valk *et al*. (2023) (Fig. 5*A*). The hair cell population present in fetal week (FW) 7.5 and 9.2 inner ear dataset is considered immature vestibular hair cells, as cochlear hair cells arise only by FW10 of fetal development (van der Valk *et al*., 2023). In the UMAP projection (Fig. 5*B*), the reprogrammed hair cells overlapped with developing fetal hair cells, while iPSCs and the other fetal inner ear cell populations formed distinct clusters separate from the reprogrammed and primary hair cells. Based on the expression pattern of 1526 genes (representing the top 100 differentially expressed genes in each cluster), the reprogrammed hair cells exhibit the highest Spearman’s correlation (0.729) with fetal hair cells, designating this cluster (among the 15 distinct cell populations in the fetal dataset) as the closest resemblance to human iHCs (Fig. 5*C*). Differential expression analysis shows that the top Gene Ontology (GO) terms in human iHCs, whether derived from the stable cell line or through retroviral reprogramming, are enriched for biological processes involved in ‘sensory perception of mechanical stimulus’ (*p* = 2.04 x 10^−8^ or 8.69 x 10^−11^) and in ‘sensory perception of sound’ (*p* = 2.82 x 10^−8^ or 1.29 x 10^−10^) (Fig. 5*D*; Supp. Fig. 9).

**Figure 5.**
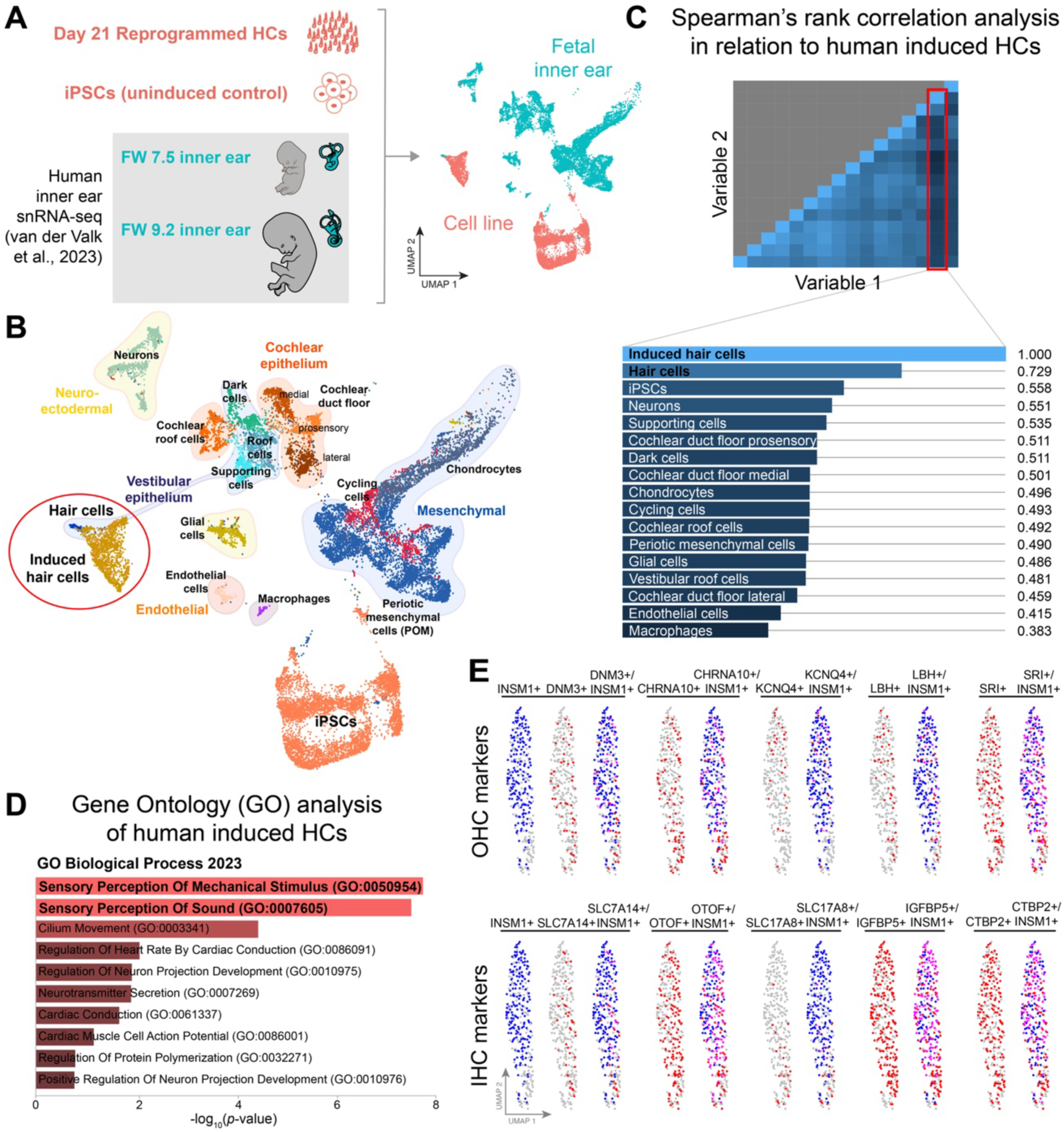
The transcriptome of human hair cell-like cells closely resembles that of fetal human hair cells and displays mixed expression patterns of different hair cell subtypes. (*A*) Single-nucleus RNA-seq of human induced HCs (cluster R5 in Fig. 3) and uninduced control iPSCs (cluster C1) were integrated with the human fetal inner ear dataset published in van der Valk *et al*. (2023) (van der Valk *et al*., 2023) and plotted using UMAP projection. (*B*) UMAP plot of integrated datasets with cell type annotations. Induced hair cells (iHCs) overlap with developing fetal hair cells. (*C*) Spearman’s rank correlation matrix for cell populations present in (*B*). iHC cluster shows the highest correlation with fetal HC cluster. (*D*) Gene Ontology (GO) analysis of the top 200 DE genes in human iHCs. GO Biological Process terms are shown, ranked by *p*- value. (*E*) Human iHCs exhibit co-expression patterns of both outer hair cell (OHC) and inner hair cell (IHC) markers. scRNA-seq profiles of cluster RV-R3 highlight this co-expression phenomenon, where cells show the simultaneous expression (colored in pink) of blue OHC subtype-specifying gene *INSM1* and red OHC markers (*DNM3*, *CHRNA10*, *KCNQ4*, *LBH*, *SRI*) or IHC markers (*SLC7A14*, *OTOF*, *SLC17A8*/*VGLUT3*, *IGFBP5*, *CTBP2*). Only cells demonstrating moderate-to-high expression levels of each gene are colored.

We further analyzed the composition of reprogrammed hair cells to determine their specific hair cell subtypes. We first examined whether known HC subtype-specifying genes are activated in human iHCs. *INSM1* and *IKZF2* (*HELIOS*) have been implicated in promoting outer hair cell (OHC) fate (Wiwatpanit *et al*., 2018; Chessum *et al*., 2018), while *TBX2* has been linked to promoting inner hair cell (IHC) fate (García-Añoveros *et al*., 2022). Analysis of scRNA-seq and snRNA-seq profiles of the reprogrammed clusters reveals expression of *INSM1*, but not *IKZF2* or *TBX2* (Supp. Fig. 10). Given that *INSM1* is known to prevent trans-differentiation into inner hair cells (Wiwatpanit *et al*., 2018), we assessed whether INSM1+ cells coincide with OHC-specific expression (such as DNM3, which is detected in the stereocilia bundles of only OHCs) (Li *et al*., 2018), to the exclusion of IHC-specific expression (such as SLC7A14, detected in the soma of only IHCs) (Li *et al*., 2018). INSM1+ cells largely coincide with markers of both OHCs (*DNM3*, *CHRNA10*, *KCNQ4*, *LBH*, *SRI*) and IHCs (*SLC7A14*, *OTOF*, *SLC17A8* [also known as *VGLUT3*], *IGFBP5*, *CTBP2*) (Fig. 5*E*), indicating a lack of differentiation between OHCs and IHCs in reprogrammed hair cells. Additionally, reprogrammed hair cells co-expressed several vestibular hair cell-associated genes, including *SOX2*, *FOXJ1*, and *ANXA4* (Supp. Figs. 11 and 12). The co-expression of established cochlear and vestibular hair cell-associated markers suggests that the reprogrammed cells have not adopted a distinct cochlear vs. vestibular identity. These findings imply that reprogrammed hair cells represent an immature or hybrid population with mixed subtype-specific expression. Overall, the data suggest that the core group SAPG can activate multiple differentiation programs specific to various hair cell types, resulting in the co-expression of OHC, IHC and vestibular HC markers within the reprogrammed cell population.

### Human induced hair cells exhibit heterogenous voltage-dependent ion currents

In our previous work, we demonstrated that SAPG-transduced mouse induced hair cells are similar in both morphological features and electrophysiological properties to primary mouse hair cells (Menendez *et al*., 2020). To analyze the voltage-gated currents and passive membrane properties of human induced hair cells, we performed whole-cell patch-clamp recordings after 7 days of reprogramming in continuous doxycycline (n = 11 cells) and compared them with uninduced control (n = 13). Using voltage-clamp, we measured the magnitude and time dependent activity of the whole-cell currents. Among the induced cells tested, some exhibited both sustained outward currents (positive currents) and transient inward (negative) currents, likely sodium currents based on their activation kinetics (Fig. 6*F*, *H*). The kinetics of the outward currents were heterogeneous: some cells displayed robust quick-activating currents that inactivated slowly over the course of the protocol (Fig. 6*F*), consistent with those observed in primary hair cells (Menendez *et al*., 2020). Others showed mixed kinetics, with currents that activated at relatively negative voltages exhibiting some inactivation, while those activating at more positive voltages were slower to activate and did not inactivate (Fig. 6*G*). Some induced cells displayed rapidly inactivating potassium currents (Fig. 6*H*), whereas a subset produced very small currents (Fig. 6*I*). The large cell-to-cell variability in the size and kinetics of the outward currents indicates heterogeneity in the ion channel composition across reprogrammed cells. In contrast, the uninduced cells were relatively homogenous exhibiting few to no currents (Fig. 6*B-E, J*). Using current-clamp (Fig. 6*L*), the human induced cells demonstrated a mean resting potential of −49 mV (± 3.1), comparable to SAPG-transduced mouse induced hair cells in monolayer culture (−50.8 mV ± 2.4; Menendez *et al*., 2020) and to previously reported values for primary hair cells (Dallos, 1985; Oliver *et al*., 2003). By contrast, the uninduced cells were more depolarized, with a mean resting potential of −24 mV (± 3.1), and struggled to maintain this potential. Together, these electrophysiological data suggest that the uninduced cells possess very few ion-selective currents, while reprogramming with SAPG yields cells with a heterogenous composition of voltage-dependent ion channels.

**Figure 6.**
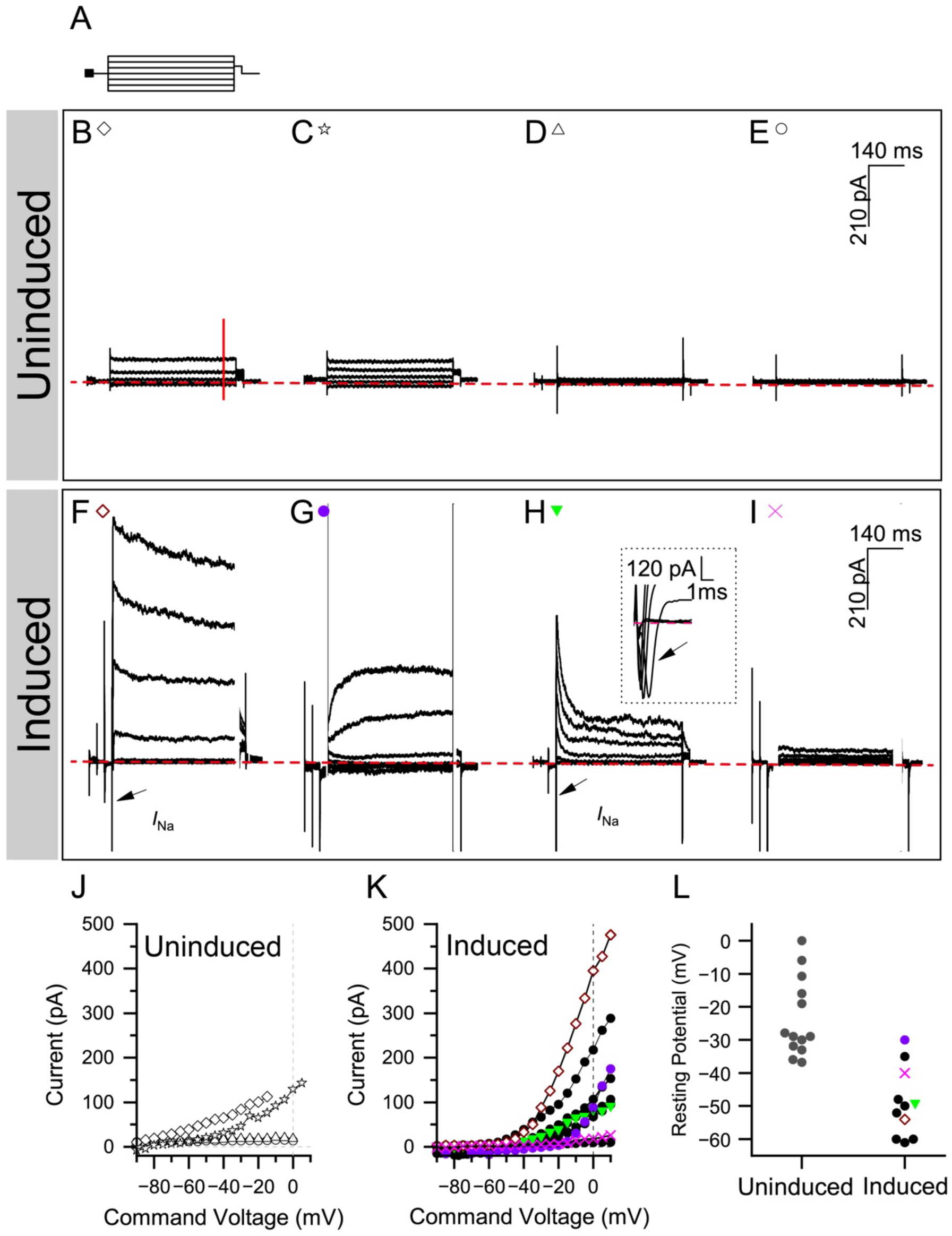
Human induced hair cells exhibit heterogenous voltage-dependent ion currents. (*A*) The voltage clamp protocol used to measure the inward and outward currents for uninduced and induced (7-day reprogrammed) cells involved voltage steps ranging from −120 to 30 mV. The voltage steps were less extreme in the uninduced cells, which uniformly had trouble holding a more negative potential. (*B*-*I)* Whole-cell currents were measured in response to a family of voltage steps (as described in *A*) in four example uninduced cells (*B*-*E*, representing the four most negative resting potentials) or in four example induced cells (*F*-*I*). A dashed red line is drawn at I=0. Sodium currents are labeled with arrows as *I*_Na_. In *H*, an inset shows *I*_Na_ at a higher resolution. (*J*-*K*) The current vs. voltage input/output functions summarize the net currents measured approximately 400 ms into the voltage step in 13 uninduced cells (*J*) and 11 induced cells (*K*). (*L*) Comparison of the resting potential of uninduced and induced cells.

## Discussion

In our efforts to address the challenges posed by the paucity and extreme inaccessibility of inner ear sensory cells for regenerative medical research and ototoxicity management, we previously identified a combination of four transcription factors, *Six1*, *Atoh1*, *Pou4f3*, and *Gfi1* (SAPG), capable of reprogramming mouse somatic cells into hair cell-like cells via direct lineage conversion using viral delivery (Menendez *et al*., 2020). We demonstrated that SAPG-transduced mouse hair cell-like cells exhibit transcriptional and epigenetic similarities to perinatal primary cochlear hair cells. Additionally, these induced hair cells recapitulate susceptibility to known ototoxins, including gentamicin (Menendez *et al*., 2020) and cisplatin (Supp. Fig. 14), validating their use for screening purposes. Building on the efficacy of the SAPG transcription factor combination, we have now adapted this approach to human cells using a virus-free induction system. Our results show that a stable iPS cell line carrying doxycycline-inducible SAPG can be reliably and effectively reprogrammed to a hair cell-like state, achieving a 19.1-fold increase in reprogramming efficiency compared to our previous retroviral-mediated strategy (Fig. 2*C*). This represents a significant improvement over the inherently low reprogramming efficiency of retroviral-mediated methods. Moreover, the inducible system eliminates the laborious and often unreliable step of converting iPSCs into secondary fibroblasts before reprogramming, which is necessitated by retroviral silencing in iPSCs. These enhancements to the translational value of direct lineage conversion improve the scalability and robustness of robotic, high-throughput screening, enabling the screening of larger drug libraries comprising tens-to-hundreds of thousands of compounds. Consequently, this approach has the potential to rapidly accelerate the screening process for identifying genes capable of inducing hair cell regeneration, as well as facilitating pre-clinical drug discovery efforts focused on ototoxicity and otoprotection.

Current retroviral- and lentiviral-mediated reprogramming strategy relies on constitutive expression of SAPG (Menendez *et al*., 2020); however, continuous expression of the master regulator ATOH1 has been shown to inhibit hair cell maturation (Liu *et al*., 2012; Liu *et al*., 2014b). The inducible system offers an advantage as the expression of the reprogramming factors can be controlled by turning them off upon withdrawal of doxycycline. Our inducible system demonstrates rapid return of the inducible SAPG factors to baseline levels after a 3- or 7- day pulse of doxycycline (Fig. 1*D*; Supp. Fig. 2). Furthermore, the stable cell line exhibits sustained expression of endogenous hair cell transcription factors and canonical hair cell markers following the pulsed treatment. Improvements to the current virus-mediated strategy will involve redesigning it to activate the master regulator ATOH1 so that its expression can be turned on to induce reprogramming and then off again, to mimic the regulation of ATOH1 as it occurs in vivo (Kelley, Woods and Montcouquiol, 2004; Kelley, 2006), as well as the switch to adeno-associated viruses (AAVs) as a superior therapeutic gene transfer platform (Zinn and Vandenberghe, 2014). Transducing supporting cells or non-sensory cells in the deafened organ of Corti for a brief period, to transiently re-enter the cell cycle to restore the correct numbers of hair cells, is a crucial characteristic needed for future gene therapy approaches to hearing loss.

Our observations indicate the SAPG reprogramming factors yield populations exhibiting mixed expression patterns characteristic of different hair cell subtypes. This results in the co-expression of markers associated with outer hair cells, inner hair cells, and vestibular hair cells within our reprogrammed population (Fig. 5*E*; Supp. Figs. 11 and 12). The concurrent expression of these markers may suggest an immature state of hair cell development. This notion is supported by previous studies showing minimal gene expression differences distinguishing inner and outer hair cells during perinatal stages (Liu *et al*., 2014a), as well between cochlear and vestibular hair cells (Burns *et al*., 2015). However, apart from *INSM1*, our reprogrammed cells fail to activate known hair cell type-specifying transcription factors, including *IKZF2* (*HELIOS*) and *TBX2* (Supp. Fig. 10), as well as *GATA3* crucial for the functional maturation of inner hair cells (Bardhan *et al*., 2019; Supp. Fig. 12). The absence of key transcription factors may explain why the SAPG combination triggers multiple differentiation programs specific to various hair cells. The lack of *IKZF2*, essential for promoting outer hair cell fate and functional maturation (Chessum *et al*., 2018), likely contributes to the failure of our mouse and human reprogrammed hair cells to express *PRESTIN* (Chessum *et al*., 2018; Menendez *et al*., 2020; Supp. Fig. 12). To address this, we plan to investigate the effects of incorporating these cochlear hair cell subtype-specifying transcription factors to the SAPG combination, which we anticipate will enhance the conversion to a cochlear hair cell fate and more advanced stage(s) of hair cell maturation. Despite not assuming a distinct cochlear vs. vestibular identity, the reprogrammed hair cells nevertheless closely resemble developing vestibular hair cells, as indicated by a Spearman’s correlation of 0.729 (Fig. 5*C*). Additionally, they demonstrate heterogenous voltage-gated ion currents (Fig. 6*F*- *I*), consistent with the mixed expression patterns of different hair cell types.

We observed upregulation of specific cochlear markers (*NR2F1/2* and *LRP2*) in a subset of reprogrammed cells, although the underlying reason(s) for this observation remain unclear. These cells form a stable cluster without a discernible trajectory (Fig. 3*B*, *D*). We hypothesize that these cells have activated the SAPG reprogramming factors but failed to fully transition to the hair cell-like state. This resembles the phenomenon of abortive reprogramming observed in previous studies on stem cell or fibroblast direct lineage conversion (Biddy *et al*., 2018; Kamimoto *et al*., 2023). We explored the role of the CMV enhancer-containing promoter driving rtTA in the inducible system, suspecting it might reduce conversion efficiency to the hair cell fate. However, prolonging incubation with iPSC medium (mTeSR) and doxycycline did not increase reprogramming factor or hair cell marker expression but was detrimental to *NR2F1/2* and *LRP2* activation (Supp. Fig. 13). The downregulation of *NR2F1/2* and *LRP2* expression in successfully reprogrammed hair cells aligns with the notion that these genes serve as cochlear markers rather than hair cell markers. Additionally, a recent study suggested that *NR2F1/2* could be involved in the temporal control of cochlear differentiation (Moore *et al*., 2023). These findings underscore the complexity of the reprogramming process and highlight the need for more comprehensive studies to elucidate the mechanisms governing inner ear cell fate determination. Nonetheless, the stable cell line shows activation of specific cochlear gene markers with inducible SAPG expression.

In reprogrammed hair cell populations derived from both human iPSCs and fibroblasts, we observed the activation of hair cell gene networks, including hair cell-specific NOTCH ligand genes (Supp. Fig. 6). This activation coincided with the downregulation of genes associated with cell cycle progression (Figs. 3 *D* and *E*; 4*A*) and *NOTCH1/2/3* receptor genes (Supp. Fig. 6). These results correspond with the characteristic postmitotic state of primary hair cells and are consistent with the documented modulation of NOTCH signaling during hair cell differentiation and maturation (Kelley, 2006). We did not detect the activation of genes associated with supporting cell identity (Supp. Fig. 4). However, previous studies have shown that overexpression of *Atoh1*, *Pou4f3*, and *Gfi1* can induce both hair cell and supporting cell networks in the mature mouse cochlea (Iyer *et al*., 2022; McGovern *et al*., 2024). This suggests that reprogramming from both iPSCs and fibroblasts may lack the additional signaling cues necessary to undergo conversion into supporting cell-like cells in response to SAPG. Indeed, iPSC- and fibroblast-derived reprogrammed hair cells failed to robustly activate some key players in the NOTCH pathway for inducing supporting cell fate, such as *DLL1*, which acts synergistically with *JAG2* to regulate hair cell differentiation and recruit neighboring cells for supporting cell development (Kiernan *et al*., 2005).

Our study underscores the potential of the inducible SAPG system in overcoming the limitations associated with retroviral- and lentiviral-mediated reprogramming of inner ear sensory cells. By achieving a more efficient, scalable, and physiologically relevant conversion of readily available somatic cell types to a hair cell-like state, our approach facilitates the high-throughput screening of genes and drugs to identify novel candidates for hair cell regeneration and otoprotection. Additionally, reprogramming from human cells is more likely to accurately reflect the effectiveness of these candidates for human use, given the potential species differences. Although the reprogrammed cells currently display mixed expression patterns characteristic of different hair cell subtypes, our future efforts will focus on refining the transcription factor combination to more precisely guide differentiation towards specific cochlear or vestibular hair cell fates. This optimization will enhance our ability to model hair cell development and maturation in vitro, ultimately advancing the development of gene and cell-based therapies for hearing restoration.

## Materials and Methods

### Generation of POU4F3-tdTomato iPSC reporter line

Human iPSCs derived from ND05280 (NINDS/Coriell; as described in Shi *et al*., 2018) were obtained from Justin Ichida. A pX330 plasmid (Addgene # 42230) expressing Cas9 and a single guide RNA (sgRNA) targeting the POU4F3 locus (GTTCGGCTGTCCACTGATTG *CGG*; PAM sequence is shown in italics) was used to introduce site-specific double-strand breaks via electroporation. A homologous recombination (HR) plasmid was co-electroporated, containing left and right homology arms (LHA and RHA), a T2A-tdTomato reporter sequence, and a puromycin resistance cassette for selection. Electroporation was performed using the Lonza nucleofector kit in the presence of 2 μM Nu7441 (STEMCELL Technologies), following the manufacturer’s instructions. After electroporation, colonies were expanded and screened by genotyping to confirm successful integration of the reporter cassette. Colonies 2 and 4 were selected for further analysis, with colony 2 showing successful amplification of both the LHA and RHA. All constructs were validated by sequencing.

### Generation of stable iPSC line carrying doxycycline-inducible *SIX1*, *ATOH1*, *POU4F3*, and *GFI1* (SAPG)

A polycistronic SAPG gBlock (GeneArt, Thermo Fisher Scientific) was designed to include coding sequences for *SIX1*, *ATOH1*, *POU4F3*, and *GFI1*, separated by self-cleaving 2A peptide sequences. The gBlock was subcloned into the CLYBL-TRE-HDR vector (referred to as CLYBL- 4F HDR; GeneArt, Thermo Fisher Scientific), which was designed with homology arms flanking the SAPG cassette for targeted integration into the CLYBL safe harbor locus. The CLYBL-TRE- HDR vector also contains a tetracycline response element (TRE) for doxycycline-inducible expression, a puromycin resistance cassette, and an mApple screenable marker.

POU4F3-tdTomato iPSCs were plated at a density of 120,000 cells per well in three wells of a 24-well plate in mTeSR medium (STEMCELL Technologies) supplemented with 10 µM ROCK Inhibitor (Y-27632, STEMCELL Technologies) and incubated for 4 hours at 37°C. Ribonucleoprotein (RNP) complexes were prepared using Alt-R® S.p. Cas9 Nuclease V3 (3 µM, IDT) in OptiMEM (Thermo Fisher Scientific) and sgRNA (3 µM, Synthego; sequence: ACTTCCTTCTATGTAAGATG *AGG*) in a 1:1 ratio, followed by a 5-minute incubation at room temperature. The RNP complex was combined with 750 ng of the CLYBL-4F HDR donor plasmid, and a mastermix of OptiMEM and Lipofectamine Stem (Thermo Fisher Scientific) was added to yield a final volume of 50 µL, followed by a 10-minute incubation at room temperature. After the 4-hour iPSC incubation, the medium was replaced with 500 µL of fresh, prewarmed mTeSR medium containing 10 µM ROCK Inhibitor. The CRISPR transfection solution (50 µL) was added dropwise to each well, and the plate was swirled gently to ensure even distribution. Cells were incubated at 32°C for 48 hours to facilitate transfection.

Following expansion, transfected cells were collected for fluorescence-activated cell sorting (FACS). mApple-positive cells were isolated and plated into single wells of a 96-well plate in mTeSR medium supplemented with CloneR (STEMCELL Technologies) to promote survival and colony formation. To generate a clonal line, positive cells were sorted again by FACS into 24 individual wells of a 96-well plate, with each well containing a single cell. Sorted cells were cultured in mTeSR medium with CloneR for 5 days, and 22 of the 24 conditions survived. These were subsequently expanded under standard mTeSR culture conditions.

After clonal expansion, the cell line was treated with 0.1 µg/µL TAT-CRE recombinase (Excellgen) in the presence of Lipofectamine Stem to excise the mApple screenable marker. Cells were FACS-sorted as mApple-negative cells and cultured in mTeSR medium with Penicillin-Streptomycin (Pen-Strep, Corning). The stable cell line was expanded and tested for karyotypic abnormalities at Children’s Hospital Los Angeles, showing an apparently normal karyotype.

### Reprogramming

POU4F3-tdTomato iPSCs carrying doxycycline-inducible SAPG were plated in mTeSR medium. Once the iPSCs reached 80% confluence, they were dissociated into single cells using 0.5 mM EDTA (Thermo Fisher Scientific) and re-plated in hair cell medium (HCM), consisting of DMEM/F- 12 + GlutaMAX (Gibco), 1% N2 supplement (Gibco), 2% B27 supplement (Gibco), 5 ng/mL EGF (Miltenyi Biotec), and 2.5 ng/mL FGF (Miltenyi Biotec), in the presence of 2 µg/mL Doxycycline monohydrate (D1822, Sigma-Aldrich). Cultures were plated with ROCK Inhibitor to promote survival for the first 24 hours, with fresh medium provided daily. For retroviral reprogramming, POU4F3-tdTomato iPSCs were converted into secondary fibroblasts as previously described in Shi *et al*. (2018) and transduced with retroviruses as previously described in Menendez *et al*. (2020).

### Immunostaining

Cells for staining were washed with phosphate-buffered saline (PBS) and fixed using 4% paraformaldehyde (PFA) in PBS for 15 min at room temperature. For permeabilization and blocking, cells were incubated in tris-buffered saline (TBS) containing 0.5% Tween-20 and 10% normal donkey serum (NDS) for 2 hours at room temperature. After blocking, the cells were washed three times with TBS for 10 min each. Cells were then incubated overnight at 4°C with the primary antibody diluted in blocking solution (TBS with 10% NDS). Following primary antibody incubation, cells were washed three times in TBS and then incubated with the appropriate Alexa Fluor 488-conjugated secondary antibody diluted in TBS for 90 min at room temperature in the dark. The DNA was stained with DAPI (1:10,000 dilution in TBS) for 10 min at room temperature.

### Western blot analysis

Cells for immunoblotting were lysed in RIPA lysis buffer (Cell Signaling Technology) supplemented with cOmplete EDTA-free Protease Inhibitor (Roche), 10 µg/mL DNAase I, and 10 mM MgCl_2_ for 30 min at 4°C. Total protein concentrations were determined using the BCA assay (Thermo Fisher Scientific). Lysates were either stored at −80°C or used immediately for subsequent analysis. Proteins were resolved on a NuPAGE 4–12% Bis-Tris Gel (Thermo Fisher Scientific) and transferred to an Immobilon-FL PVDF membrane (Merck Millipore). Membranes were incubated overnight at 4°C with the primary antibody diluted in blocking solution (5% BSA in TBS) containing 0.5% Tween-20. After incubation, membranes were washed five times in TBST (TBS with 0.1% Tween-20) at room temperature and subsequently incubated with the appropriate Alexa Fluor 680-conjugated secondary antibody diluted in blocking solution containing 0.2% Tween-20 and 0.1% SDS for 1 hour at room temperature in the dark. The membranes were imaged using an Odyssey CLx imaging system (LI-COR).

### Antibodies

Anti-MyosinVI (Proteus Bioscience, catalog# 25-6791; 1:500 for immunostaining, 1:5,000 for Western)

Anti-MyosinVIIa (Proteus Bioscience, catalog# 25-6790; 1:1000 for immunostaining, 1:10,000 for Western)

Anti-Pou4f3 (Santa Cruz Biotechnology, catalog# sc-81980; 1:1000 for immunostaining, 1:10,000 for Western)

Anti-HA tag (Abcam, catalog# ab9110; 1:4,000) Anti-Nanog (Abcam, catalog# ab173368; 1:2,000)

Anti-Pou5f1/Oct4 (Invitrogen, catalog# 701756; 1:500) Anti-β-Actin (Sigma-Aldrich, catalog# A2228; 1:10,000)

Anti-rabbit Alexa Fluor 488 (Invitrogen, catalog# A21206; 1:500)

Anti-mouse Alexa Fluor 488 (Invitrogen, catalog# A21202; 1:500)

Anti-rabbit Alexa Fluor Plus 680 (Invitrogen, catalog# A32802; 1:10,000) Anti-mouse Alexa Fluor Plus 680 (Invitrogen, catalog# A10038; 1:10,000)

### Imaging

Immunostaining images of adherent cell cultures were acquired on a Leica SP8-X confocal microscope. For quantifying reprogramming efficiency in adherent cultures, images were captured at 10x magnification with the Molecular Devices ImageExpress and processed using CellProfiler, as described below.

### iHC detection and counting method

Automated cell counting was performed using CellProfiler 4.2.1, with thresholds set for size, intensity, and roundness of the fluorescence signal. Imaging for this analysis was conducted at 10x magnification. For each image, a size filter was applied to remove objects that were either too large or too small. The remaining objects were detected through adaptive thresholding using the Minimum Cross-Entropy method. The presence of each fluorescence marker was quantified as total number of cells per well and as a percentage of the total number of DAPI+ cells per well. To ensure the reliability of automated counting, a comparison was conducted between manual and automated counts across 8 images, with no significant difference observed (*p* = 0.85).

### RNA isolation and RT-qPCR

RNA was isolated using the RNeasy Micro Kit (Qiagen). For reverse transcription, RNA was converted to cDNA using the SuperScript IV VILO Master Mix with ezDNase Enzyme (Thermo Fisher Scientific). Quantitative PCR (qPCR) was performed using the Luna Universal qPCR Master Mix (New England Biolabs) according to the manufacturer’s protocol, on a ViiA 7 Real-Time PCR System with a 96-well block (Thermo Fisher Scientific). Because the polycistronic SAPG induction cassette in the stable cell line was designed with synonymous codon changes relative to endogenous SAPG, transgene-specific and endogenous-specific primers were used to distinguish transgenic SAPG from endogenous SAPG. These codon differences do not alter the encoded protein sequence. The primer sequences used in this study are as follows:

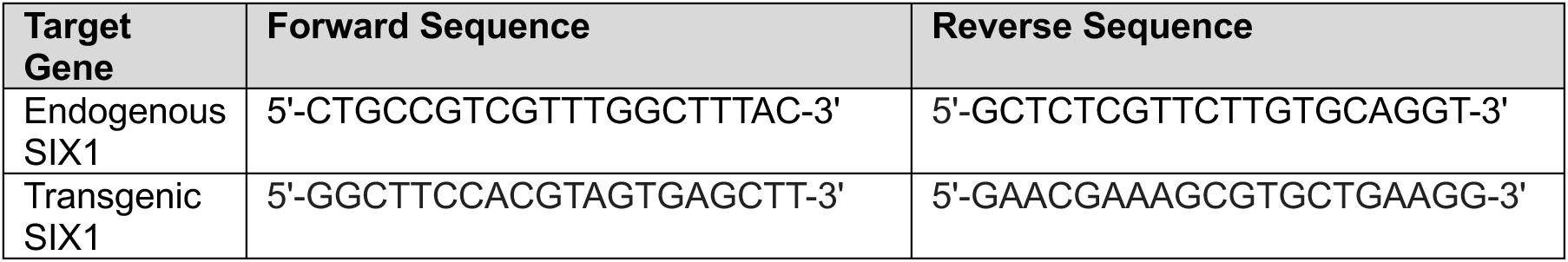
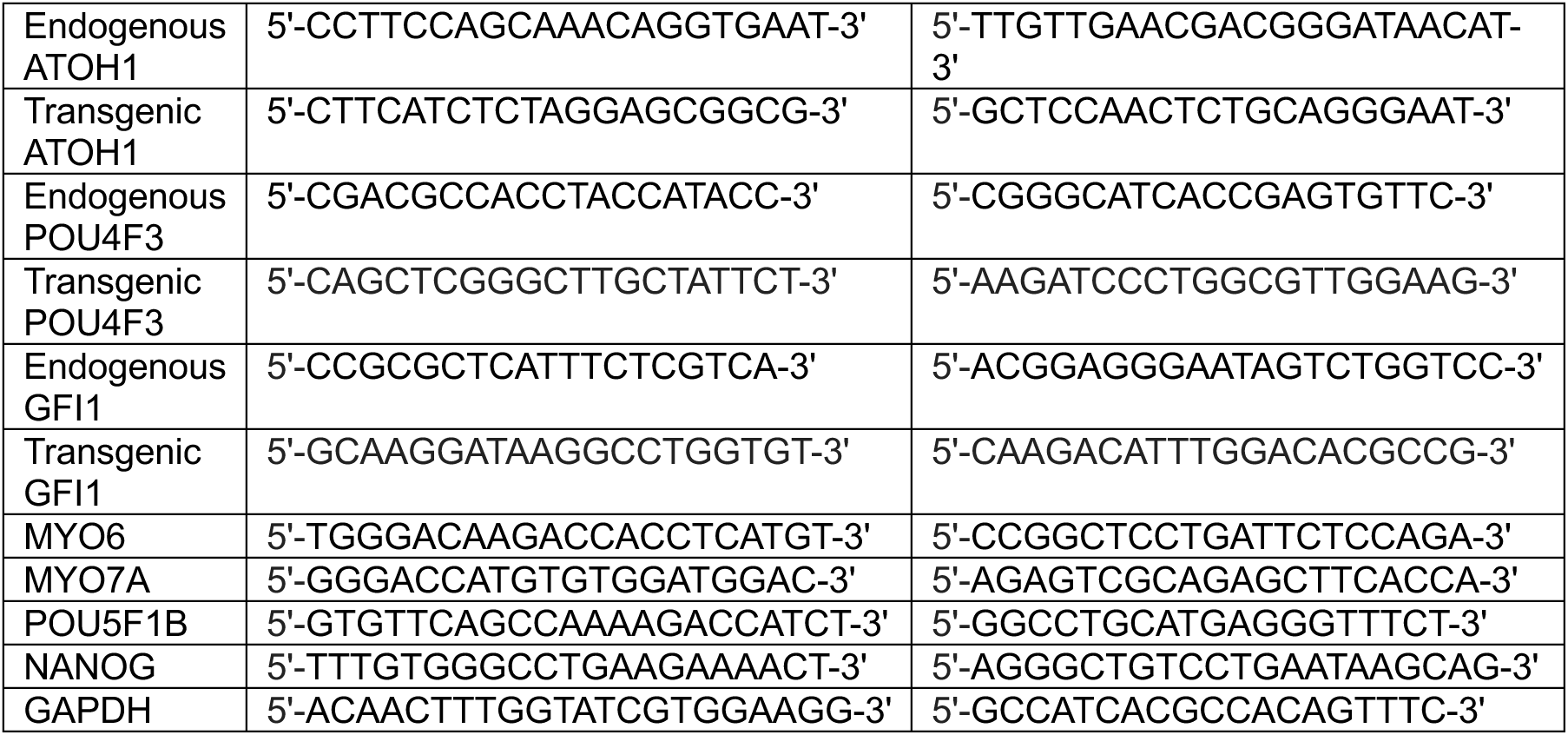

### Single-cell and single-nucleus RNA sequencing

For Day 21 retroviral-reprogrammed iHCs, POU4F3-tdTomato cells transduced with SAPG were FACS-sorted into tdTomato^+^ and tdTomato^−^ populations. A combined population consisting of 16,945 tdTomato^+^ cells and 16,945 tdTomato^−^ cells was processed using the Chromium Next GEM Single Cell 3’ Reagent Kits (CG000204; 10x Genomics), following the manufacturer’s instructions, yielding a final recovery of 6,676 single cells. For Day 21 iHCs reprogrammed from the stable cell line, cells treated with doxycycline continuously for 21 days and uninduced control were processed using the Nuclei Isolation for Single Cell Multiome ATAC + Gene Expression Sequencing protocol (CG000365; 10x Genomics). Briefly, cells were EDTA-dissociated and resuspended in 1 mL PBS + 1% BSA. Nuclei were filtered using a Falcon 5 mL round bottom polystyrene test tube with a 40 μm cell strainer snap cap, then centrifuged at 500 x g for 5 minutes at 4°C. The supernatant was decanted, and nuclei were resuspended in 1 mL of PBS + 2% BSA and incubated for 5 minutes at 4°C. Nuclei were centrifuged again at 500 x g for 5 min at 4°C, and the pellet was resuspended in 100 μL of 0.1x Lysis Buffer by pipetting up and down. Nuclei were incubated on ice for 2 minutes, followed by the addition of 1 mL Wash Buffer. The suspension was mixed by pipetting. A sample was taken at this stage to quantify cell concentration. The remaining nuclei suspension was centrifuged at 500 x g for 5 minutes at 4°C, and the nuclei pellet was resuspended in the appropriate volume of Diluted Nuclei Buffer to achieve approximately 2000 nuclei/μL for input into the Single Cell Multiome ATAC + Gene Expression protocol (CG000338; 10x Genomics). For the Day 21 induced sample, 5,152 quality nuclei data points were recovered. For the uninduced control, 6,181 quality nuclei data points were recovered.

### scRNAseq and snRNAseq data processing

Raw sequencing data in FASTQ format were processed using the 10x Genomic Cell Ranger 4.0 or Cell Ranger ARC 2.0.0 pipeline (http://support.10xgenomics.com/). The sequencing reads were aligned to either the human reference GRCh38 genome (POU4F3-tdTomato) or a custom GRCh38 genome reference that incorporates the polycistronic SAPG cassette sequence (POU4F3-tdTomato-CLYBL-4F). The filtered gene expression matrices were used as input into Seurat v4.1.1 for standard quality control, preprocessing, and filtering of low-quality data points. This resulted in 6,676 cells for Day 21 retroviral-reprogrammed sample, 5,028 nuclei for Day 21 doxycycline-induced sample, and 6,048 nuclei for uninduced control. Following data preprocessing, normalization and scaling were performed using SCTransform, with highly variable features identified for downstream analysis. Principal component analysis (PCA) was computed based on the identified variable genes. Datasets from the Day 21 doxycycline-induced sample and the uninduced control were integrated using SCT-based normalization for account for differences across conditions. These integrated datasets were visualized and clustered using Uniform Manifold Approximation and Projection (UMAP). Quantitative and specific gene expression patterns were visualized using R packages ggplot2 and viridis. RNA velocity analysis was performed using Velocyto.R, applying gene-relative velocity estimates and embedding velocity vectors onto the UMAP projection. For integration with the human fetal inner ear snRNAseq dataset published in van der Valk *et al*. (2023), the fetal week 9.2 and week 7.5 datasets (GEO accession number GSE213796) were combined. Normalization, scaling, dimensional reduction and clustering were performed as described in van der Valk *et al*. (2023). Subsetted C1 and R5 (representing iPSCs and reprogrammed hair cells) dataset was integrated with the fetal inner ear dataset using the same workflow. These integrated datasets were clustered using UMAP at a resolution of 0.9, and clusters were assigned cell type identities based on known cell markers, as described in van der Valk *et al*. (2023).

Sequencing data have been uploaded to the GEO database (https://www.ncbi.nlm.nih.gov/geo) under accession number GSE351733. All scripts, processed Seurat objects, and metadata annotations are available in the GEO repository. The deposited resources include workflows for normalization, integration, clustering, and visualization to ensure reproducibility.

### GO and GSEA analyses

Gene ontology analysis was performed and visualized using Enrichr (Chen *et al*., 2013). Gene Set Enrichment Analysis was performed using GSEA v4.3.2 (Subramanian *et al*., 2005) with 1000 permutations and classic (non-weighted) statistic. The ranked list metric for GSEA was based the formula log_2_(fold change)*−log_10_(adjusted *p*-value).

### Electrophysiology

Electrophysiological recording was conducted as previously described in Menendez *et al*. (2020). Briefly, whole cell patch clamping was performed on induced (7-day reprogrammed) and uninduced POU4F3-tdTomato iPSCs carrying doxycycline-inducible SAPG. Preparations were viewed at X630 using a Zeiss Axios Examiner D1 microscope fitted with Zeiss W Plan-Aprochromat optics. Signals were driven, recorded, and amplified with an Multiclamp 700B amplifier, Digidata 1440 board and pClamp 10.7 software (pClamp, RRID:<u>SCR_011323</u>). Recording and cleaning pipettes were fabricated using filamented borosilicate glass. Pipettes were fired polished to yield an access resistance between 4–8 MΩ. Each recording pipette was covered in a layer of parafilm to reduce pipette capacitance. Recording pipettes were filled with one of two internal solutions, a standard internal solution and a perforated patch internal solution. The contents of the standard internal solution are (in mM): 135 KCl, 3.5 MgCl_2_, 3 Na_2_ATP, 5 HEPES, 5 EGTA, 0.1 CaCl_2_, 0.1 Li-GTP, and titrated with 1 M KOH to a pH of 7.35 and an osmolarity of 300 mmol/kg. The contents of the perforated-patch solution are (in mM): 75 K_2_SO_4_, 25 KCl, 5 MgCl_2_, 5 HEPES, 5 EGTA, 0.1 CaCl_2_, and titrated with 1 M KOH to a pH of 7.4 and an osmolality of 260–270 mmol/kg. Amphotericin B (240 µg/mL final concentration; Sigma-Aldrich) was dissolved in DMSO and added to the perforated-patch solution on the day of recording. This allowed passage of small monovalent ions while preventing larger molecules from dialyzing. The voltage clamp protocol was performed by holding the cell at −60 mV followed by a stimulus of voltage steps (−120 to +70 mV, by intervals of 10 mV). Uninduced cells did not often tolerate the −60 mV holding potential; as a result in a subset of recordings we held those cells at a more modest −25 mV with greater success in cell survival. To improve the ability to record from the more fragile uninduced cells, we employed two approaches: the conventional rupture patch technique and a gentler perforated-patch method, in which the antibiotic amphotericin is used to create membrane pores, enabling electrical continuity.

Analysis of the data was performed using a combination of pClamp (pClamp, RRID:<u>SCR_011323</u>), and Origin Pro (OriginPro, RRID:<u>SCR_015636</u>). pClamp software was used to gather and quantify raw data from electrophysiological recordings.

### Statistics

Sample numbers, experimental repeats and statistical tests used are indicated in figure legends. Unless otherwise stated, data are presented as mean ± SEM of at least three biological replicates. For comparisons across multiple time points, statistical significance was assessed by one-way ANOVA followed by Tukey’s honestly significant difference (HSD) test for multiple pairwise comparisons. Adjusted *p* <.05 was considered statistically significant. All statistical analyses are provided in Supplementary file 1.

## Acknowledgments

We thank Welly Makmura and Juan Llamas for laboratory maintenance and John Duc Nguyen, Shun-Yang Cheng, and Kari Koppitch for bioinformatics support. We also acknowledge Valerie Hsiao for technical assistance, Francis James for maintaining the bioinformatics server, and Mickey Huang and the Choi Family Therapeutic Screening for high-content imaging and data transfer.

Additionally, we acknowledge both past and present lab members—Ksenia Gnedeva, Talon Trecek, Xizi Wang, Haoze Yu, Tuo Shi, Leah Kim, and Tuba Ege—for their valuable scientific discussions.

## Additional information

### Funding

| Funder | Grant reference number | Principal Investigator |
| --- | --- | --- |
| Department of Defense (DoD) | W81XWH-22-1-1009 | Neil Segil<br>Andrew McMahon<br>John Oghalai |
| California Institute for<br>Regenerative Medicine<br>(CIRM) | DISC2-10773 | Neil Segil<br>Justin Ichida |
| National Institutes of Health<br>(NIH) | R01DC015829 | Neil Segil<br>Andrew McMahon |
| National Institutes of Health<br>(NIH) | R01DC014832 | Andrew Groves |
| Keck School of Medicine of<br>USC |  | Neil Segil |
| The funders played no role in study design, data collection, interpretation, or the decision to submit the work for publication. |  |  |

### Author contributions

Robert N. Rainey, Conceptualization, Data curation, Software, Formal analysis, Funding acquisition, Investigation, Methodology, Project administration, Writing - original draft; Sam D. Houman, Conceptualization, Formal analysis, Investigation, Methodology, Writing - review and editing; Louise Menendez, Conceptualization, Formal analysis, Investigation, Methodology, Writing - review and editing; Ryan Chang, Investigation; Litao Tao, Data curation; Formal analysis, Writing - review and editing; Helena Bugacov, Investigation, Methodology, Writing - review and editing; Andrew P. McMahon, Supervision, Project administration, Writing - review and editing; Radha Kalluri, Conceptualization, Resources, Data curation, Formal analysis, Supervision, Investigation, Methodology, Project administration, Writing - review and editing; John S. Oghalai, Supervision, Project administration, Writing - review and editing; Andrew K. Groves, Conceptualization, Supervision, Methodology, Writing - review and editing; Neil Segil, Conceptualization, Resources, Formal analysis, Supervision, Funding acquisition, Investigation, Methodology, Project administration.

## Figures and Tables

**Supplementary Figure 1.**
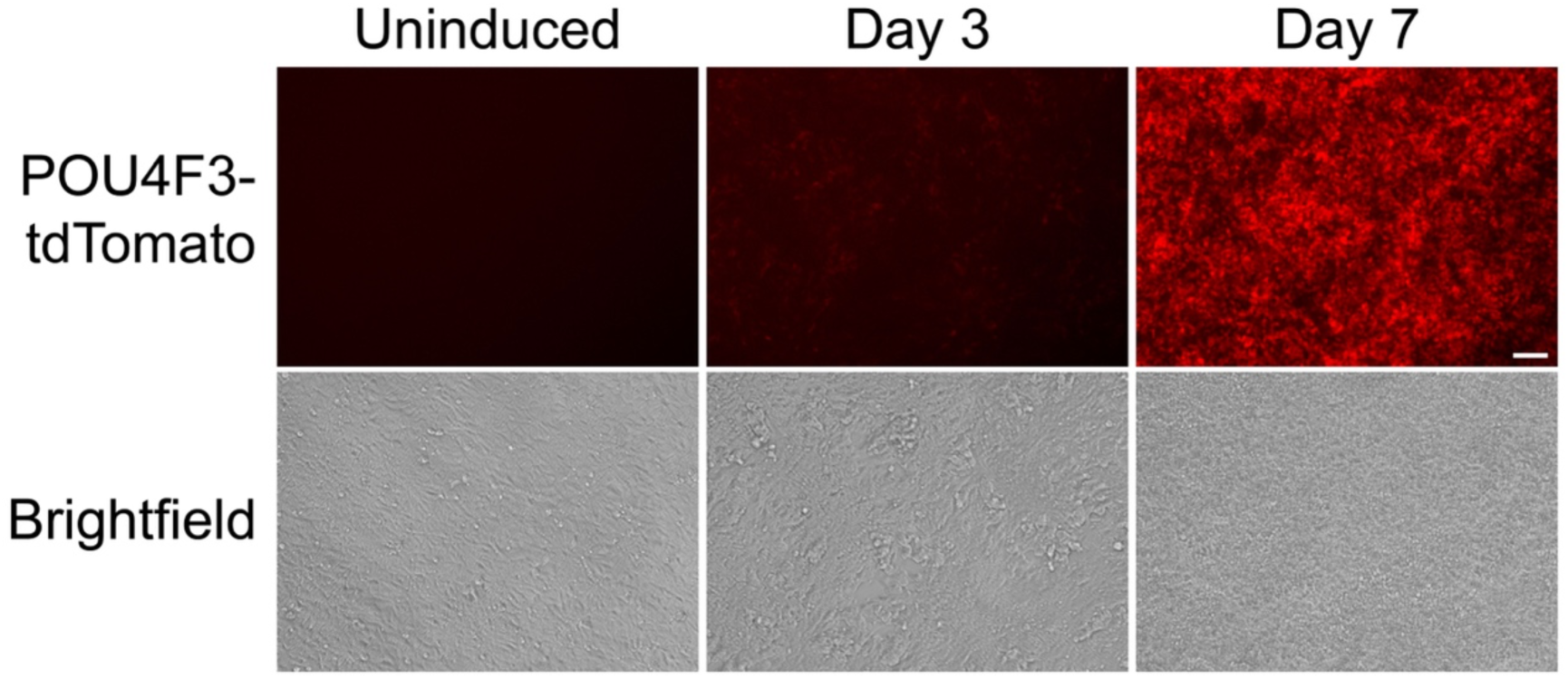
The *POU4F3*-tdTomato hair cell reporter becomes visible after ∼3 days of reprogramming. Representative images were acquired at 0-, 3- or 7-days post-doxycycline treatment. Scale bar represents 200 µm.

**Supplementary Figure 2.**
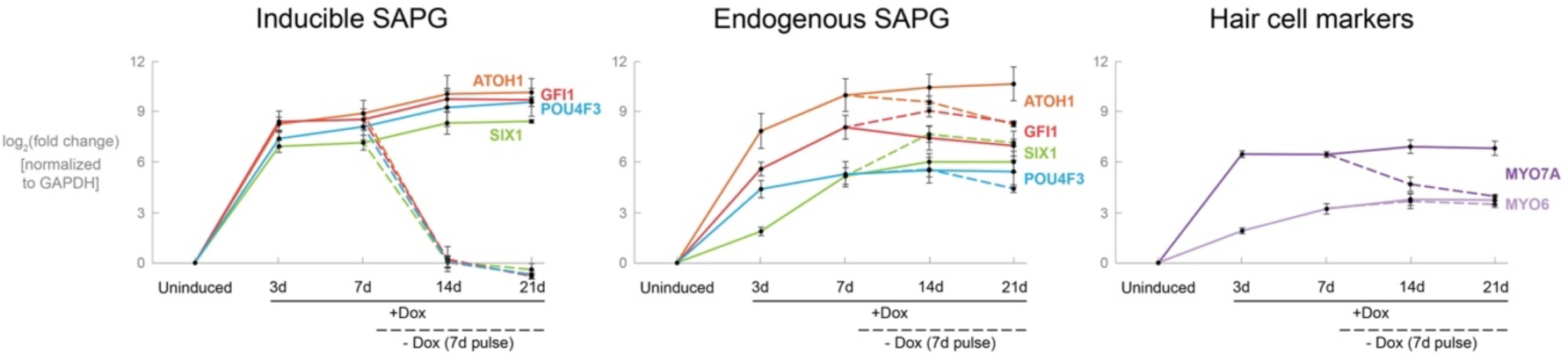
RT-qPCR analysis of the cell line treated with continuous doxycycline or a 7-day pulse of doxycycline over the indicated time points. Dashed lines indicate post-doxycycline removal. Values are normalized to GAPDH, and a ratio is calculated by dividing the uninduced control (0 hour). Fold-change values are log_2_-transformed. Error bars indicate SEM. *N* = 3 biological replicates. Statistical analyses are provided in Supplementary file 1.

**Supplementary Figure 3.**
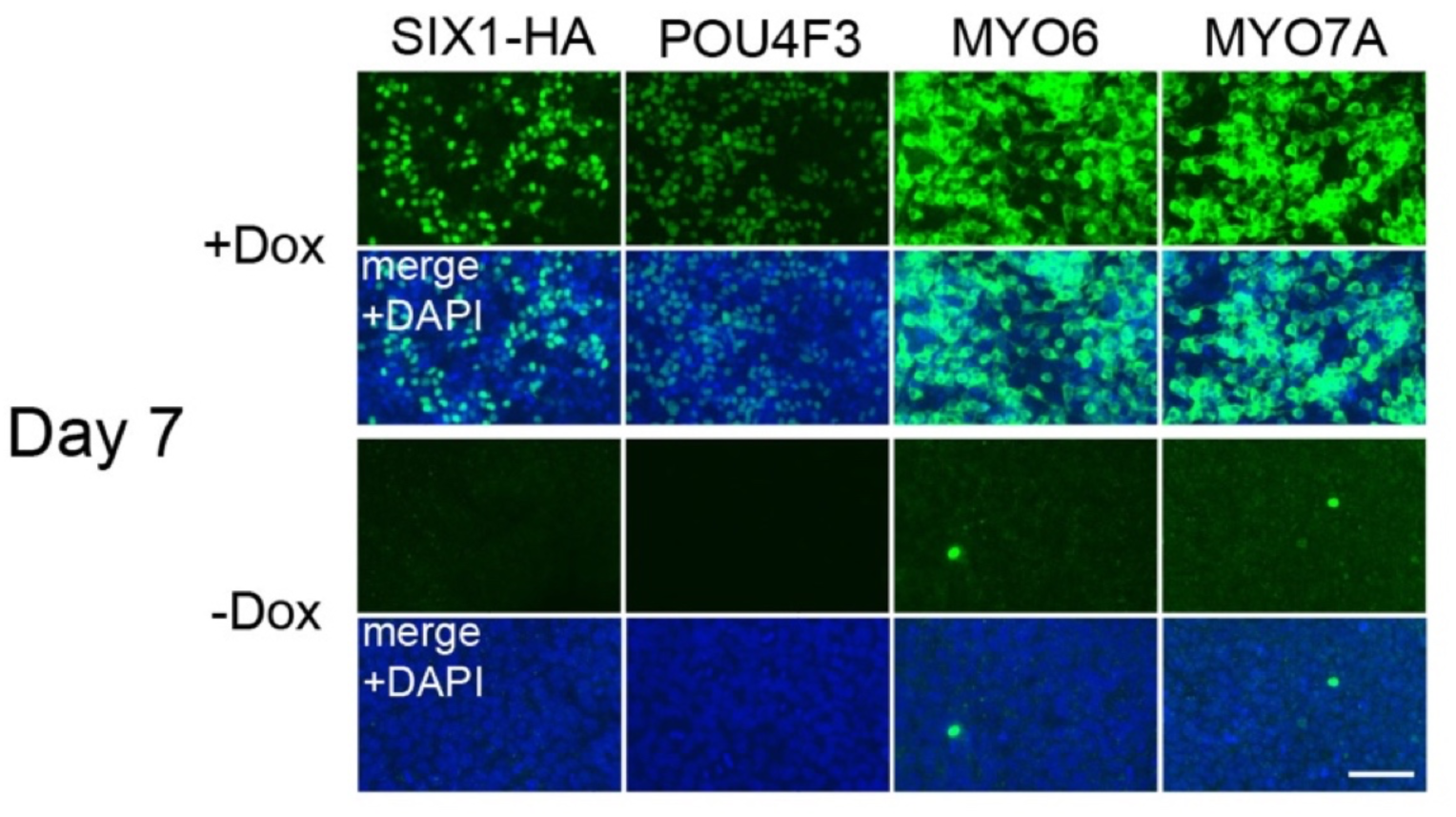
Representative pictures of cultures treated with continuous doxycycline for 7 days (+Dox) or left untreated (−Dox).

**Supplementary Figure 4.**
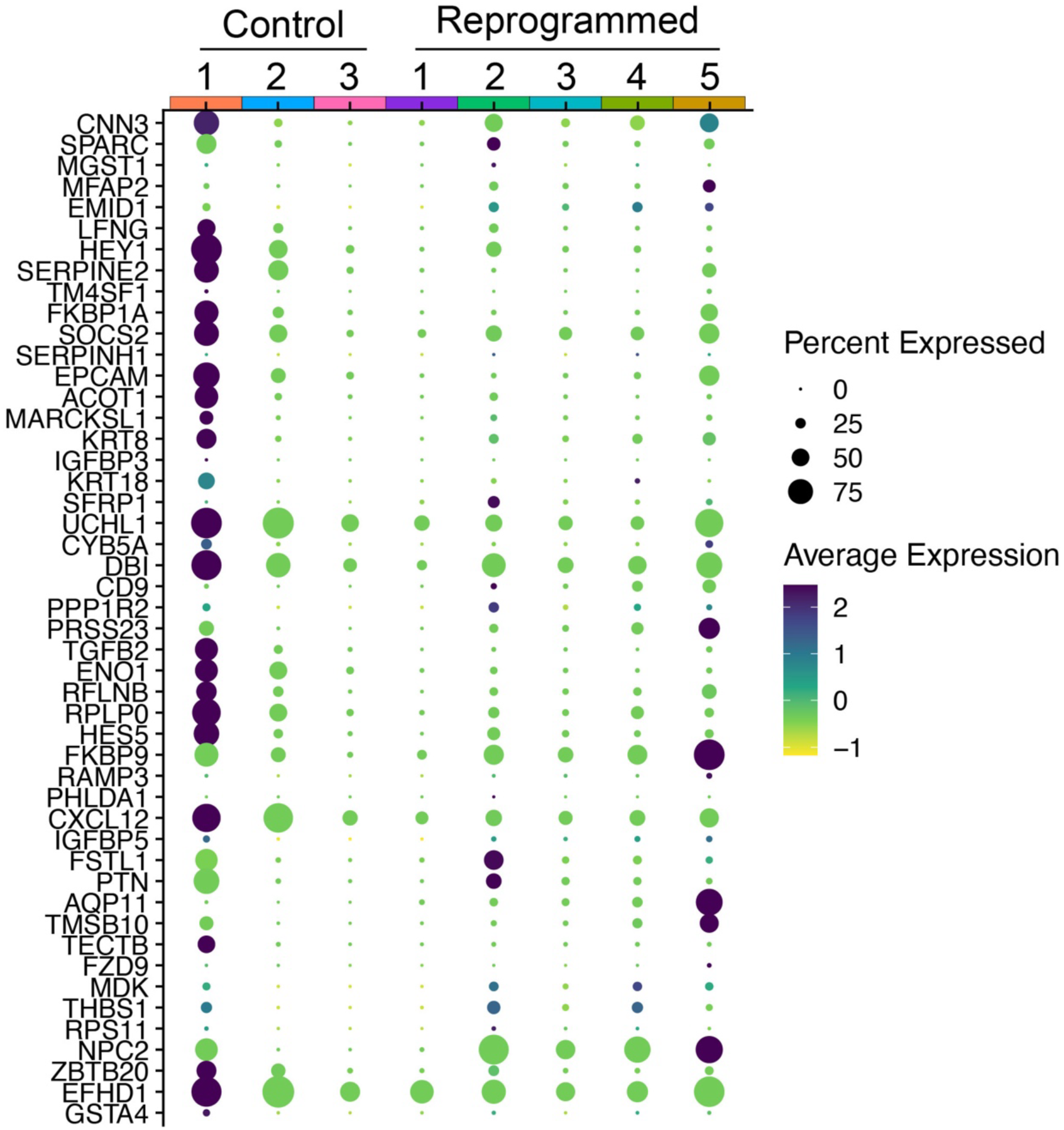
Dot plot of the top 50 DE genes in P1 supporting cells, as previously identified by Kolla *et al*. (2020), shown across our control (C1–C3) and reprogrammed (R1–R5) clusters.

**Supplementary Figure 5.**
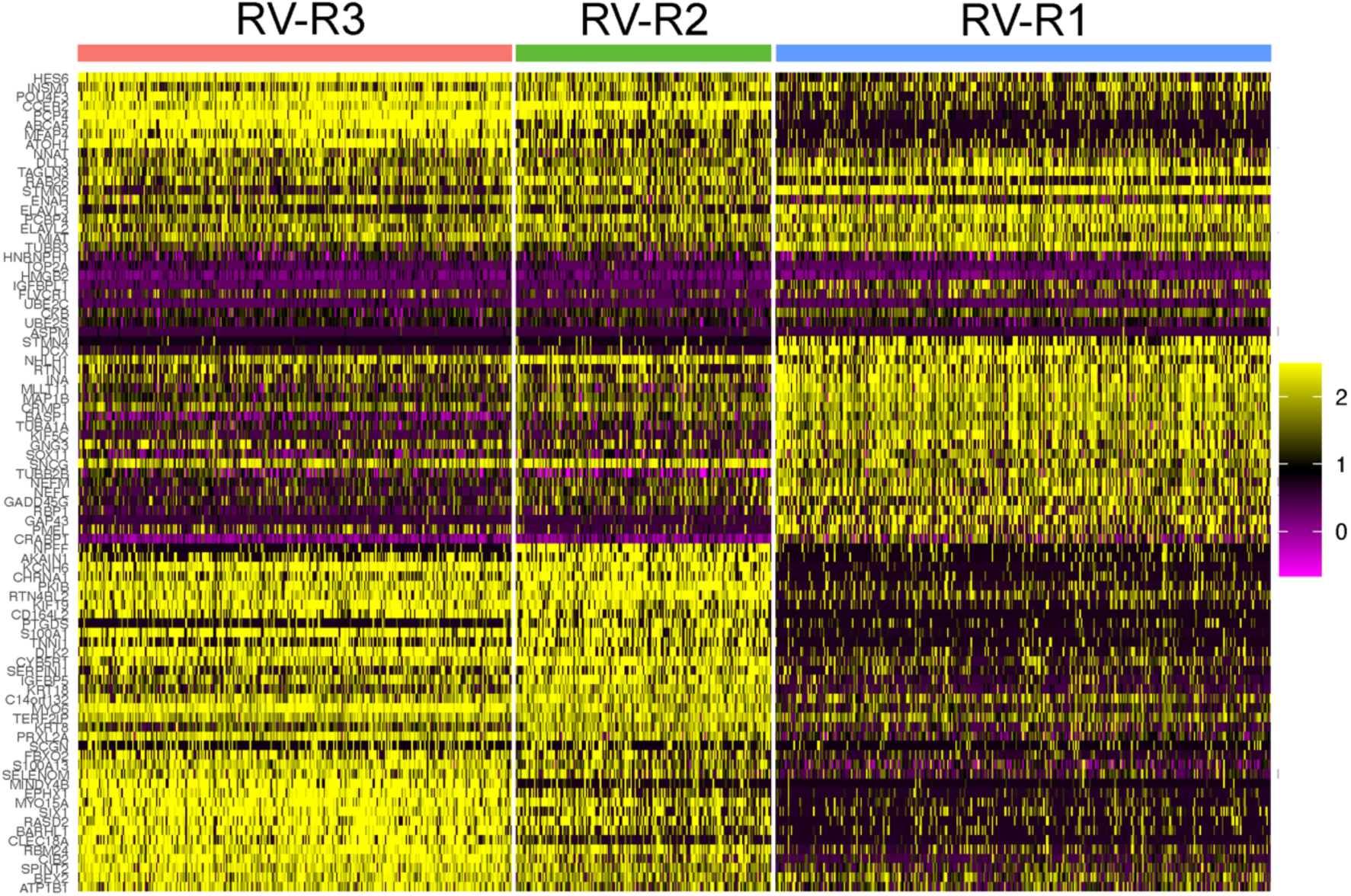
Expanded version of Figure 4B displaying the full set of differentially expressed genes.

**Supplementary Figure 6.**
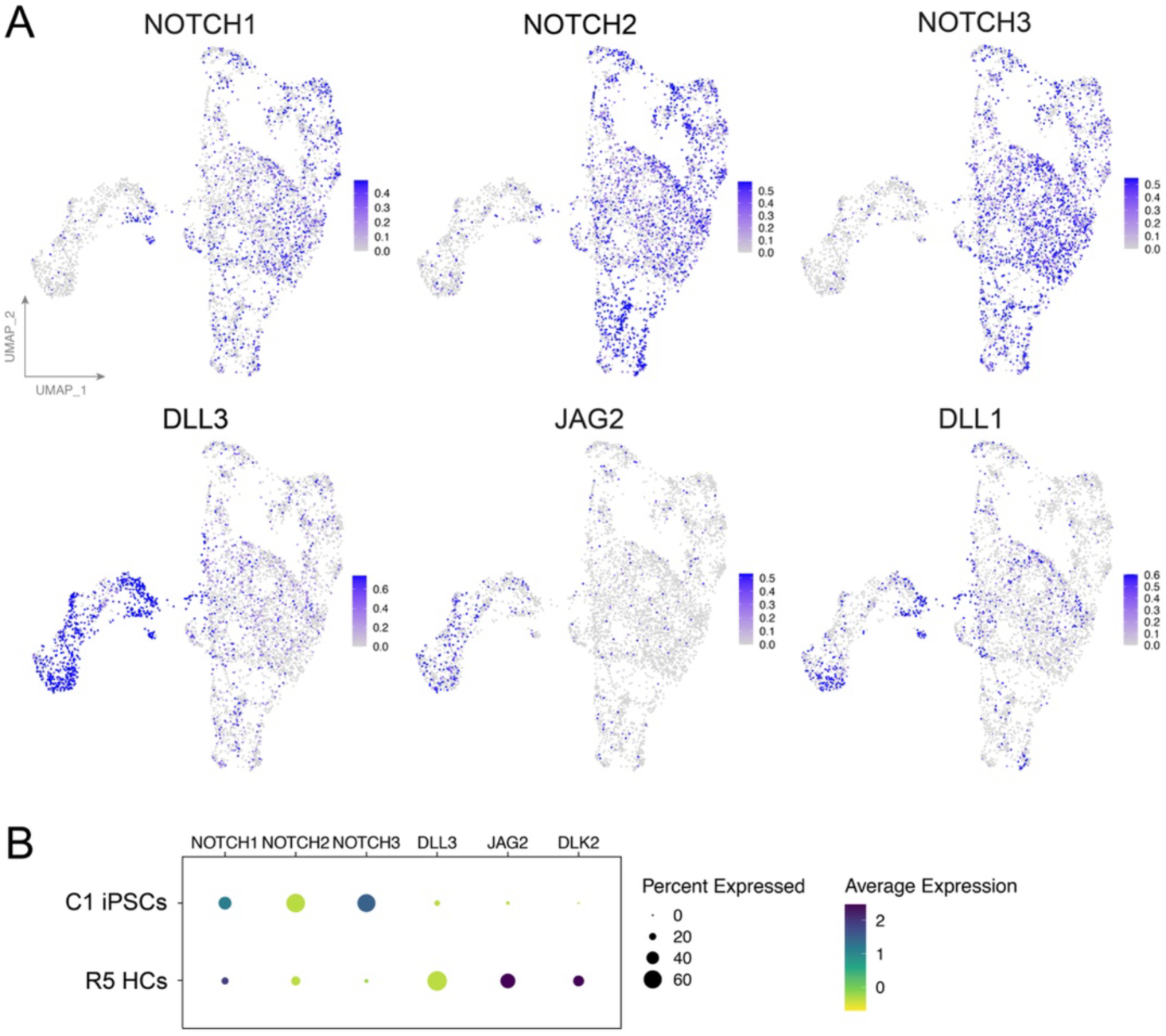
HC-like clusters show reduced expression of *NOTCH1/2/3* and increased expression of hair cell-specific NOTCH ligand genes *DLL3*, *JAG2*, and *DLK2* relative to control clusters. (*A*) UMAP projection of Day 21 reprogrammed RV-R1–3 HCs and residual fibroblasts. (B) Dot plot of Day 21 reprogrammed R5 HCs and C1 iPSCs.

**Supplementary Figure 7.**
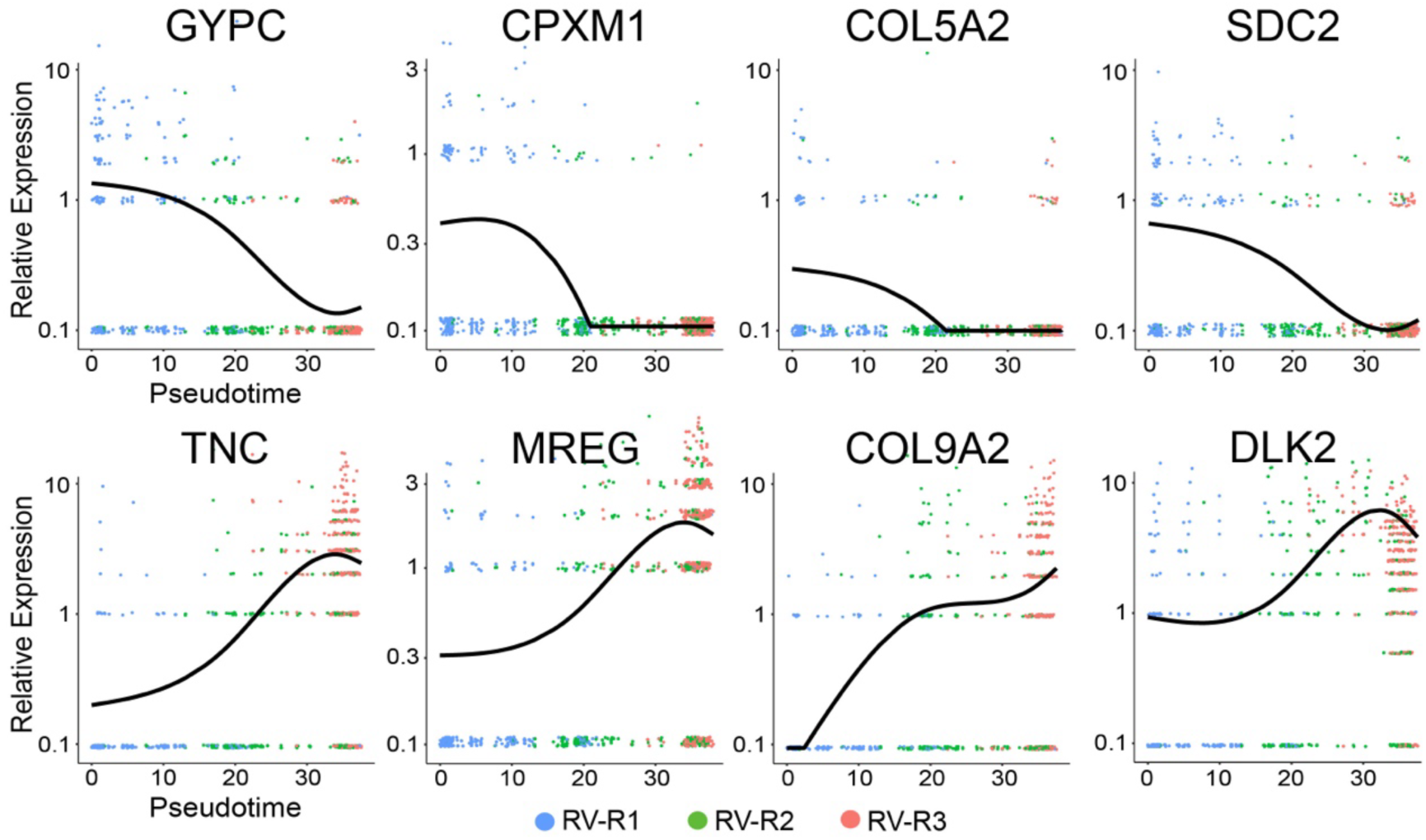
Scatterplots showing relative gene expression levels based on Monocle (pseudotime). Cells are colored by hair cell-like clusters 1–3 (RV-R1–3). Top: expression of fibroblast-enriched gene examples (*GYPC*, *CPXM1*, *COL5A2*, *SDC2*) over pseudotime. Bottom: expression of hair cell-enriched gene examples (*TNC*, *MREG*, *COL9A2*, *DLK2*).

**Supplementary Figure 8.**
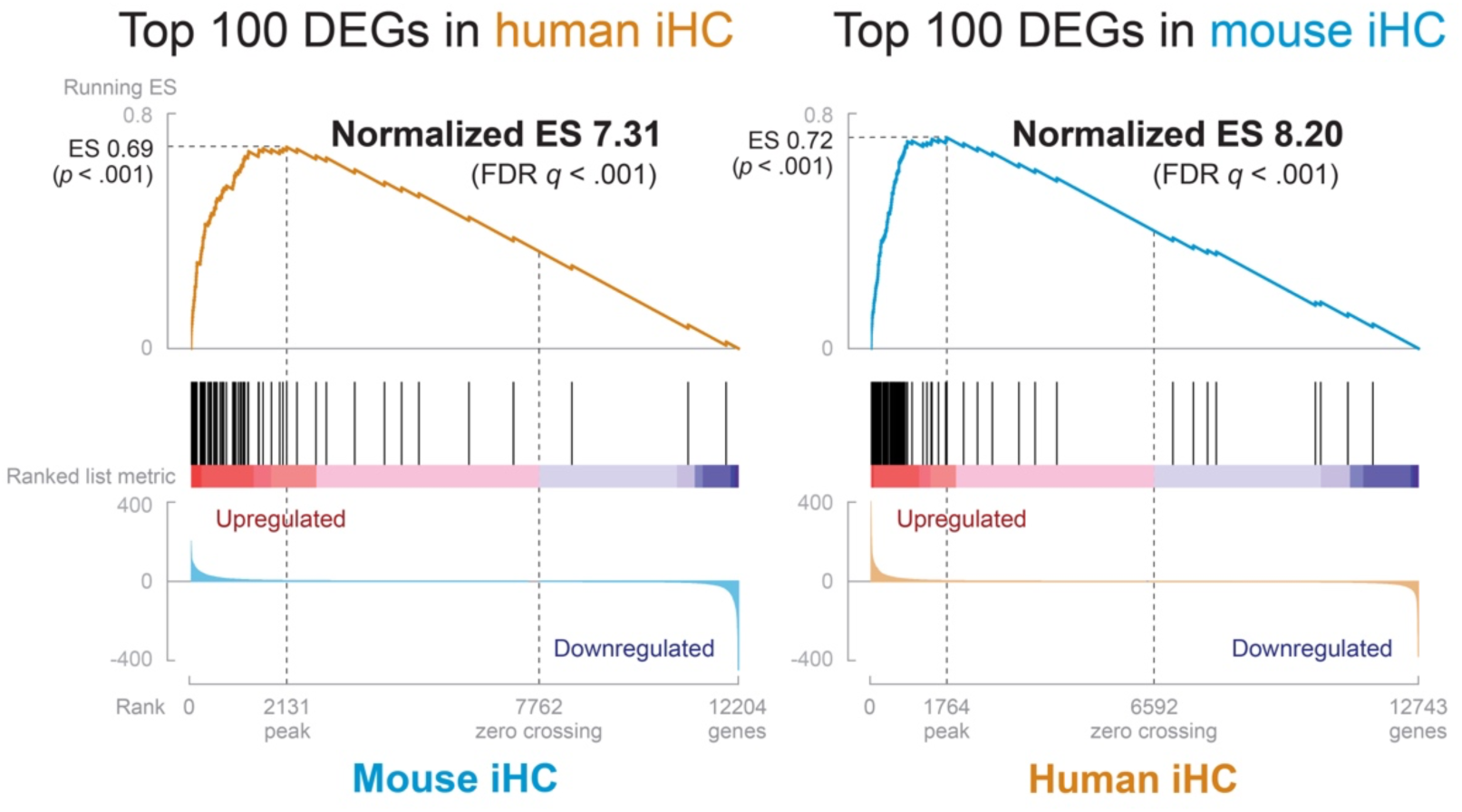
Human and mouse iHCs show highly significant enrichment of each other’s top DEGs. GSEA was used to analyze the top 100 DEGs in human iHC against mouse iHC, and the top 100 DEGs in mouse iHC against human iHC. FDR (false discovery rate) of < 25% is considered significant.

**Supplementary Figure 9.**
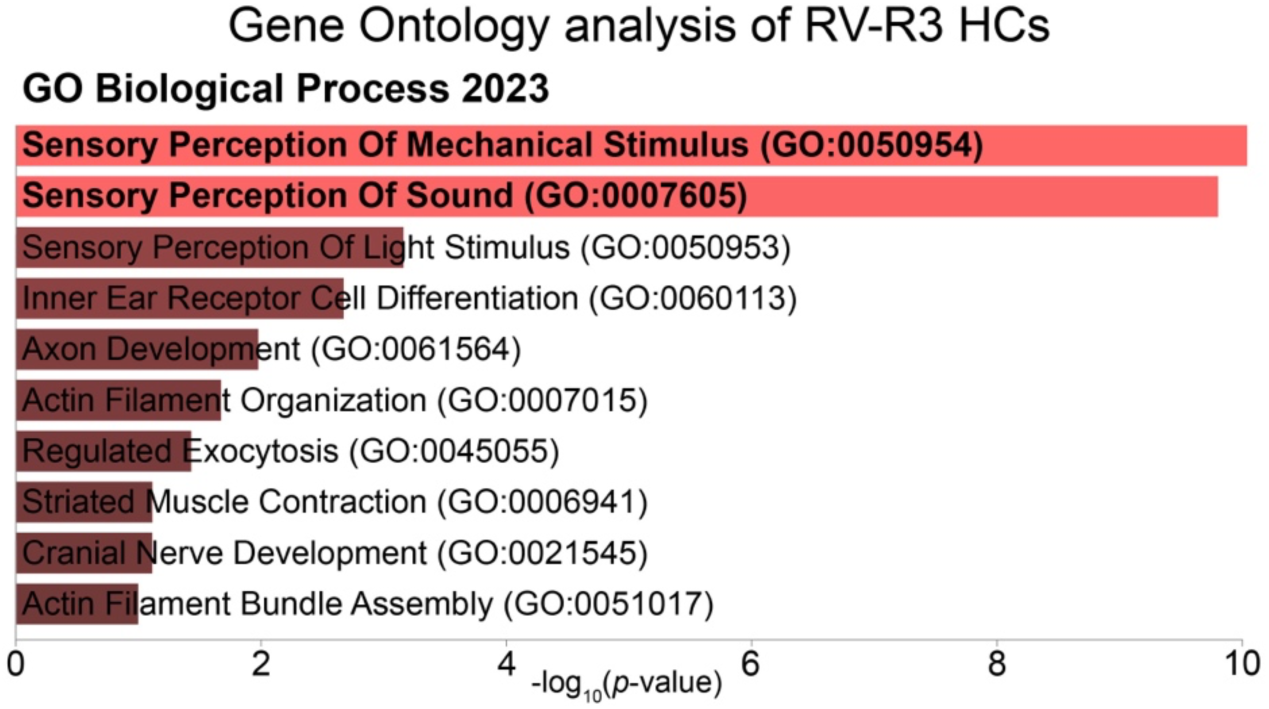
Gene Ontology (GO) analysis of the top 200 DE genes in RV-R3. GO Biological Process terms are shown, ranked by *p*-value.

**Supplementary Figure 10.**
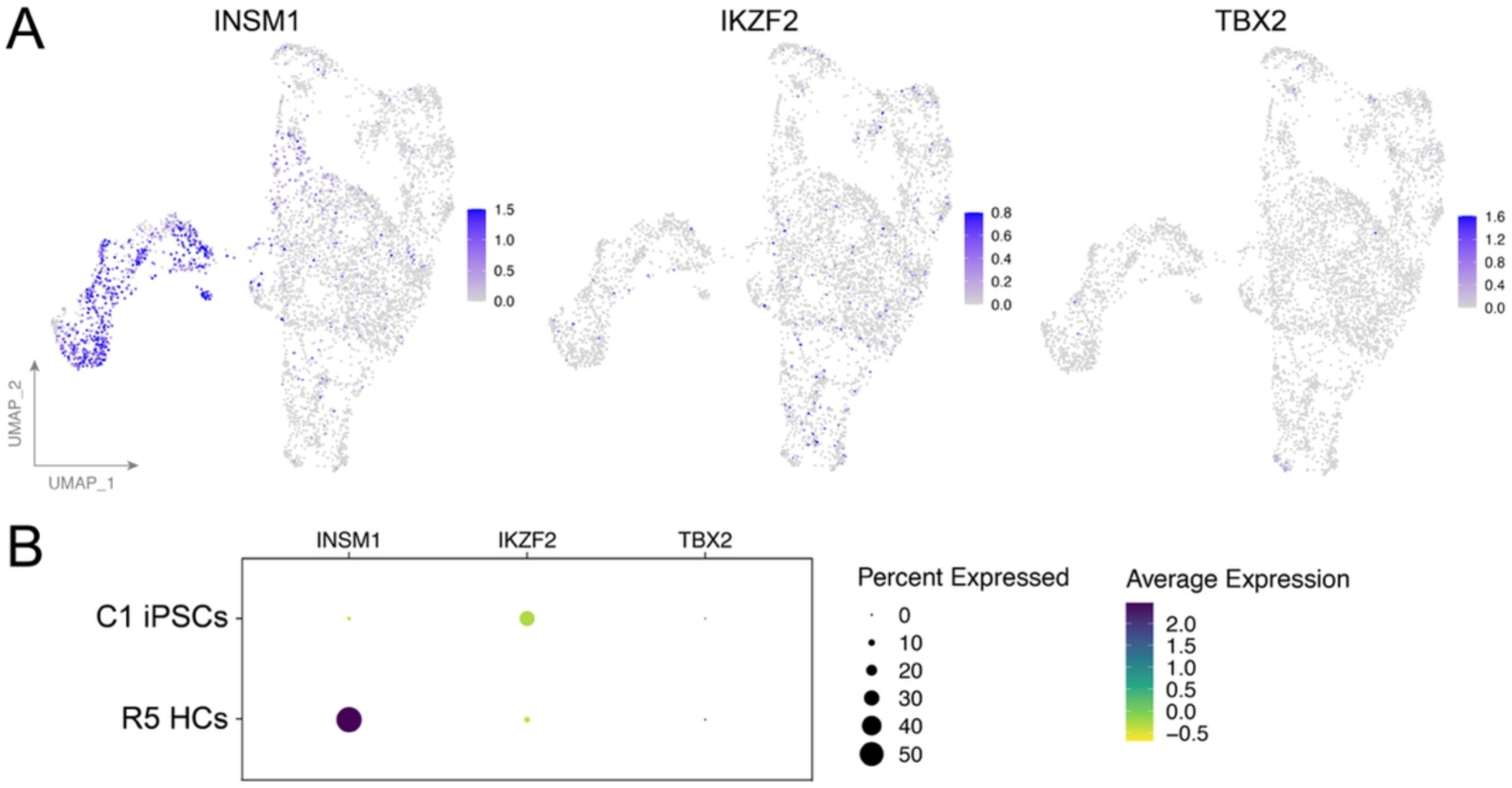
*INSM1*, but not *IKZF2* (*HELIOS*) or *TBX2*, is expressed in HC-like clusters. (*A*) UMAP projection of Day 21 reprogrammed RV-R1–3 HCs and residual fibroblasts. (B) Dot plot of Day 21 reprogrammed R5 HCs and C1 iPSCs.

**Supplementary Figure 11.**
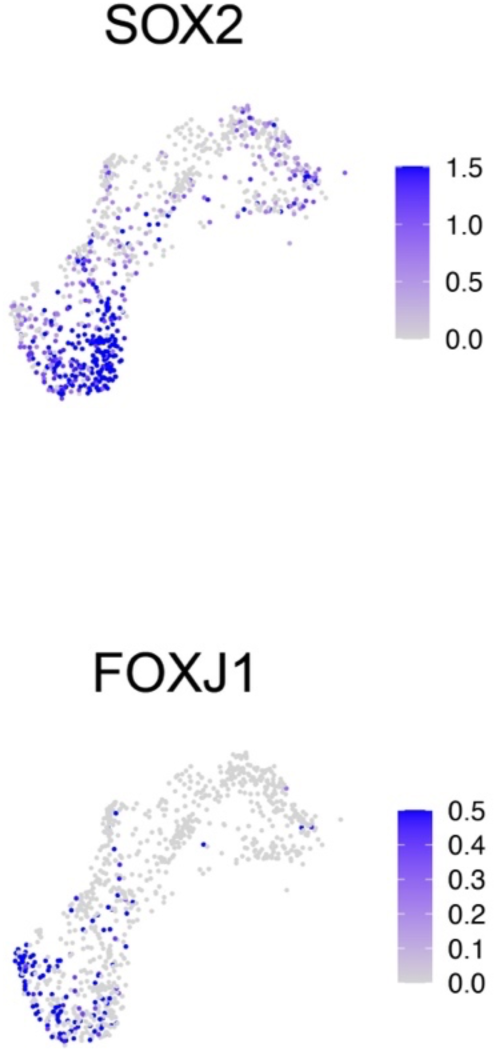
RV-R3 HCs show expression of the vestibular hair cell-associated genes *SOX2* and *FOXJ1*. SOX2, required for maintaining Type II vestibular HC identity, marks HC progenitors of both cochlear and vestibular nature but is absent from neonatal cochlear HCs. FOXJ1, involved in ciliogenesis, is expressed in vestibular HCs and neonatal cochlear HCs but not mature cochlear HCs.

**Supplementary Figure 12.**
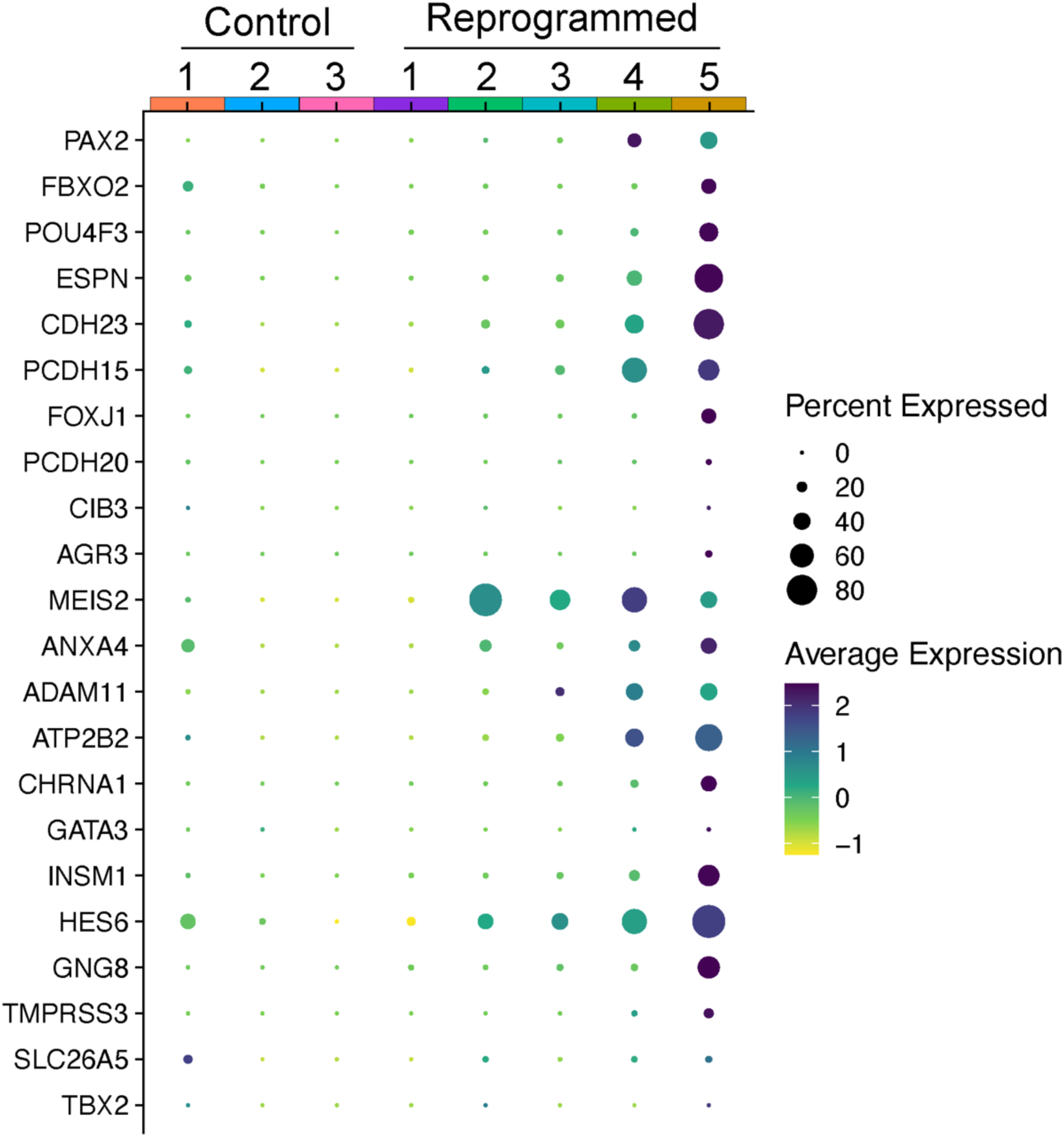
Dot plot showing expression of additional otic/hair cell-associated genes (*PAX2*, *FBXO2*, *POU4F3*, *ESPN*, *CDH23*, *PCDH15, ATP2B2,* and *CHRNA1*) and genes reported to be enriched or associated with vestibular hair cells (*FOXJ1*, *PCDH20*, *CIB3*, *AGR3*, *MEIS2*, *ANXA4*, and *ADAM11*) and cochlear hair cells (*GATA3*, *INSM1*, *HES6*, *GNG8*, *TMPRSS3*, *SLC26A5*, and *TBX2*) across C1–C3 and R1–R5 clusters.

**Supplementary Figure 13.**
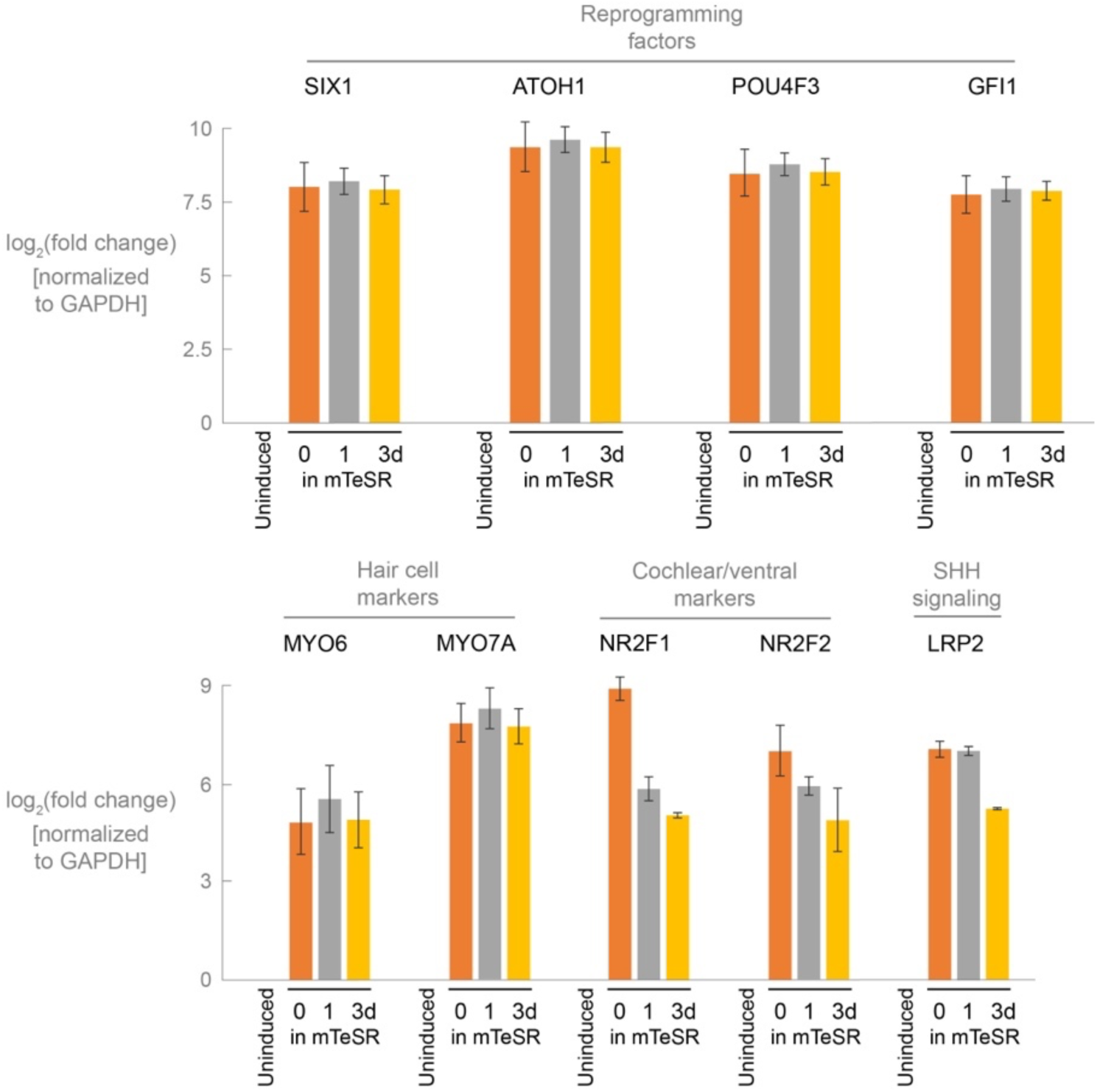
RT-qPCR analysis of the cell line treated with continuous doxycycline for 7 days. Cells were treated with doxycycline in mTeSR for 0, 1, or 3 days and moved to hair cell media + doxycycline for the balance of the 7-day reprogramming. Values are normalized to GAPDH, and a ratio is calculated by dividing the uninduced control (0 hour). Fold-change values are log_2_-transformed. Error bars indicate SEM. Statistical analyses are provided in Supplementary file 1.

**Supplementary Figure 14.**
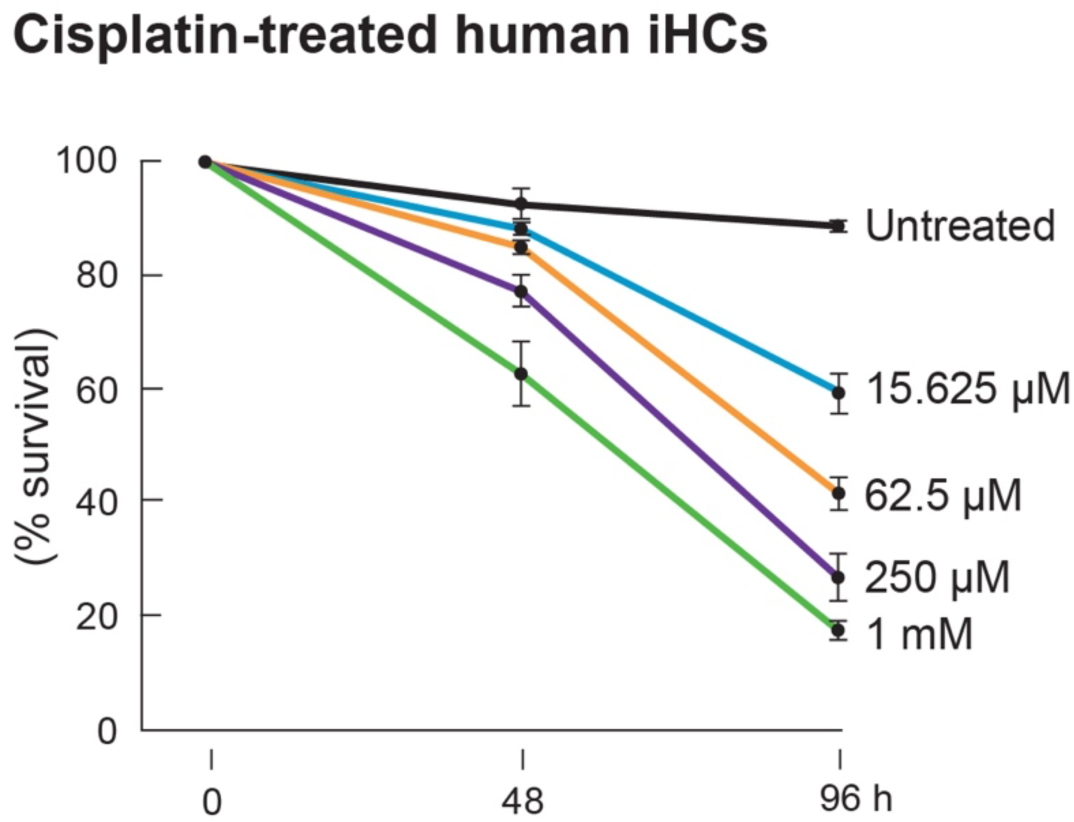
Human induced hair cells (iHCs) are sensitive to cisplatin in a dose-dependent manner. Cisplatin ototoxicity profile in human iHCs. Error bars indicate SEM. The ototoxicity assay was performed as previously described in Menendez *et al*. (2020). Statistical significance was assessed separately at 48 and 96 h by one-way ANOVA followed by Dunnett’s multiple-comparisons test comparing each cisplatin concentration with the untreated control. *p* < .05 was considered statistically significant. Statistical analyses are provided in Supplementary file 1.

## References

Alonso, M. B. D., Hernandez, I. L., Angel de la Fuente, M., Garcia-Sancho, J., Giraldez, F. and Schimmang, T. (2018) ‘Transcription factor induced conversion of human fibroblasts towards the hair cell lineage’, PloS one, 13(7), pp. e0200210–e0200210.

Babos, K. N., Galloway, K. E., Kisler, K., Zitting, M., Li, Y., Shi, Y., Quintino, B., Chow, R. H., Zlokovic, B. V. and Ichida, J. K. (2019) ‘Mitigating Antagonism between Transcription and Proliferation Allows Near-Deterministic Cellular Reprogramming’, Cell stem cell, 25(4), pp. 486–500.e9.

Bardhan, T., Jeng, J. Y., Waldmann, M., Ceriani, F., Johnson, S. L., Olt, J., Rüttiger, L., Marcotti, W. and Holley, M. C. (2019) ‘Gata3 is required for the functional maturation of inner hair cells and their innervation in the mouse cochlea’, The Journal of physiology, 597(13), pp. 3389–3406.

Biddy, B. A., Kong, W., Kamimoto, K., Guo, C., Waye, S. E., Sun, T. and Morris, S. A. (2018) ‘Single-cell mapping of lineage and identity in direct reprogramming’, Nature (London), 564(7735), pp. 219–224.

Bodmer, D. (2008) ‘Protection, regeneration and replacement of hair cells in the cochlea: Implications for the future treatment of sensorineural hearing loss’, Schweizerische Medizinische Wochenschrift, 138(47-48), pp. 708–712.

Burns, J. C., Kelly, M. C., Hoa, M., Morell, R. J. and Kelley, M. W. (2015) ‘Single-cell RNA-Seq resolves cellular complexity in sensory organs from the neonatal inner ear’, Nature communications, 6(1), pp. 8557–8557.

Carey, B. W., Markoulaki, S., Hanna, J., Saha, K., Gao, Q., Mitalipova, M. and Jaenisch, R. (2009) ‘Reprogramming of murine and human somatic cells using a single polycistronic vector’, Proceedings of the National Academy of Sciences - PNAS, 106(1), pp. 157–162.

Chan, E. M., Ratanasirintrawoot, S., Park, I.-H., Manos, P. D., Loh, Y.-H., Huo, H., Miller, J. D., Hartung, O., Rho, J., Ince, T. A., Daley, G. Q. and Schlaeger, T. M. (2009) ‘Live cell imaging distinguishes bona fide human iPS cells from partially reprogrammed cells’, Nature biotechnology, 27(11), pp. 1033–U100.

Chardin, S. and Romand, R. (1995) ‘REGENERATION AND MAMMALIAN AUDITORY HAIR-CELLS’, Science (American Association for the Advancement of Science), 267(5198), pp. 707–709.

Chen, P. and Segil, N. (1999) ‘p27(Kip1) links cell proliferation to morphogenesis in the developing organ of Corti’, Development (Cambridge), 126(8), pp. 1581–1590.

Chen, Y., Gu, Y., Li, Y., Li, G.-L., Chai, R., Li, W. and Li, H. (2021) ‘Generation of mature and functional hair cells by co-expression of Gfi1, Pou4f3, and Atoh1 in the postnatal mouse cochlea’, Cell reports (Cambridge), 35(3), pp. 109016–109016.

Cheng, A. G., Cunningham, L. L. and Rubel, E. W. (2005) ‘Mechanisms of hair cell death and protection’, Current opinion in otolaryngology & head and neck surgery, 13(6), pp. 343–348.

Chessum, L., Matern, M. S., Kelly, M. C., Johnson, S. L., Ogawa, Y., Milon, B., McMurray, M., Driver, E. C., Parker, A., Song, Y., Codner, G., Esapa, C. T., Prescott, J., Trent, G., Wells, S., Dragich, A. K., Frolenkov, G. I., Kelley, M. W., Marcotti, W., Brown, S. D. M., Elkon, R., Bowl, M. R. and Hertzano, R. (2018) ‘Helios is a key transcriptional regulator of outer hair cell maturation’, Nature (London), 563(7733), pp. 696–724.

Dallos, P. (1985) ‘Membrane potential and response changes in mammalian cochlear hair cells during intracellular recording’, The Journal of neuroscience, 5(6), pp. 1609–1615.

Faridi, R., Yousaf, R., Gu, S., Inagaki, S., Turriff, A. E., Pelstring, K., Guan, B., Naik, A., Griffith, A. J., Adadey, S. M., Aboagye, E. T., Awandare, G. A., Morell, R. J., Tsilou, E., Noyes, A. G., Sulmonte, L. A. G., Wonkam, A., Schrauwen, I., Leal, S. M., Azaiez, H., Brewer, C. C., Riazuddin, S., Hufnagel, R. B., Hoa, M., Zein, W. M., Dios, J. K. and Friedman, T. B. (2023) ‘Variants of LRP2, encoding a multifunctional cell-surface endocytic receptor, associated with hearing loss and retinal dystrophy’, Clinical genetics, 103(6), pp. 699–703.

García-Añoveros, J., Clancy, J. C., Foo, C. Z., García-Gómez, I., Zhou, Y., Homma, K., Cheatham, M. A. and Duggan, A. (2022) ‘Tbx2 is a master regulator of inner versus outer hair cell differentiation’, Nature (London), 605(7909), pp. 298–303.

Groves, A. K. (2010) ‘The challenge of hair cell regeneration’, Experimental Biology and Medicine, 235(4), pp. 434–446.

Géléoc, G. S. G. and Holt, J. R. (2014) ‘Sound Strategies for Hearing Restoration’, Science (American Association for the Advancement of Science), 344(6184), pp. 596–596.

Iyer, A. A. and Groves, A. K. (2021) ‘Transcription Factor Reprogramming in the Inner Ear: Turning on Cell Fate Switches to Regenerate Sensory Hair Cells’, Frontiers in cellular neuroscience, 15, pp. 660748–660748.

Iyer, A. A., Hosamani, I., Nguyen, J. D., Cai, T., Singh, S., McGovern, M. M., Beyer, L., Zhang, H., Jen, H. I., Yousaf, R., Birol, O., Sun, J. J., Ray, R. S., Raphael, Y., Segil, N. and Groves, A. K. (2022) ‘Cellular reprogramming with ATOH1, GFI1, and POU4F3 implicate epigenetic changes and cell-cell signaling as obstacles to hair cell regeneration in mature mammals’, eLife, 11.

Jeong, M., O’Reilly, M., Kirkwood, N. K., Al-Aama, J., Lako, M., Kros, C. J. and Armstrong, L. (2018) ‘Generating inner ear organoids containing putative cochlear hair cells from human pluripotent stem cells’, Cell death & disease, 9(9), pp. 922–13.

Kamimoto, K., Adil, M. T., Jindal, K., Hoffmann, C. M., Kong, W., Yang, X. and Morris, S. (2023) ‘Gene regulatory network reconfiguration in direct lineage reprogramming’, Stem Cell Reports, (7735), pp. 97–112.

Kelley, M. W. (2006) ‘Regulation of cell fate in the sensory epithelia of the inner ear’, Nature reviews. Neuroscience, 7(11), pp. 837–849.

Kelley, M. W., Woods, C. and Montcouquiol, M. (2004) ‘Math1 regulates development of the sensory epithelium in the mammalian cochlea’, Nature neuroscience, 7(12), pp. 1310–1318.

Kiernan, A. E., Cordes, R., Kopan, R., Gossler, A. and Gridley, T. (2005) ‘The Notch ligands DLL1 and JAG2 act synergistically to regulate hair cell development in the mammalian inner ear’, Development (Cambridge), 132(19), pp. 4353–4362.

Koehler, K. R., Mikosz, A. M., Molosh, A. I., Patel, D. and Hashino, E. (2013) ‘Generation of inner ear sensory epithelia from pluripotent stem cells in 3D culture’, Nature, 500(7461), pp. 217–21.

Koehler, K. R., Nie, J., Longworth-Mills, E., Liu, X. P., Lee, J., Holt, J. R. and Hashino, E. (2017) ‘Generation of inner ear organoids containing functional hair cells from human pluripotent stem cells’, Nat Biotechnol, 35(6), pp. 583–589.

Langer, T., am Zehnhoff-Dinnesen, A., Radtke, S., Meitert, J. and Zolk, O. (2013) ‘Understanding platinum-induced ototoxicity’, Trends in pharmacological sciences (Regular ed.), 34(8), pp. 458–469.

Lee, Y.-S., Liu, F. and Segil, N. (2006) ‘A morphogenetic wave of p27Kip1 transcription directs cell cycle exit during organ of Corti development’, Development (Cambridge), 133(15), pp. 2817–2826.

Li, Y., Liu, H., Giffen, K. P., Chen, L., Beisel, K. W. and He, D. Z. Z. (2018) ‘Transcriptomes of cochlear inner and outer hair cells from adult mice’, Scientific data, 5(1), pp. 180199–180199.

Liu, H., Pecka, J. L., Zhang, Q., Soukup, G. A., Beisel, K. W. and He, D. Z. Z. (2014a) ‘Characterization of transcriptomes of cochlear inner and outer hair cells’, The Journal of neuroscience, 34(33), pp. 11085–11095.

Liu, Z., Dearman, J. A., Cox, B. C., Walters, B. J., Zhang, L., Ayrault, O., Zindy, F., Gan, L., Roussel, M. F. and Zuo, J. (2012) ‘Age-dependent in vivo conversion of mouse cochlear pillar and Deiters’ cells to immature hair cells by Atoh1 ectopic expression’, The Journal of neuroscience, 32(19), pp. 6600–6610.

Liu, Z., Fang, J., Dearman, J., Zhang, L. and Zuo, J. (2014b) ‘In vivo generation of immature inner hair cells in neonatal mouse cochleae by ectopic Atoh1 expression’, PloS one, 9(2), pp. e89377–e89377.

Lo, C.-A., Greben, A. W. and Chen, B. E. (2017) ‘Generating stable cell lines with quantifiable protein production using CRISPR/Cas9-mediated knock-in’, BioTechniques, 62(4), pp. 165–174.

Lowenheim, H., Furness, D. N., Kil, J. and Zinn, C. (1999) ‘Gene disruption of p27Kip1 allows cell proliferation in the postnatal and adult organ of Corti’, Proceedings of the National Academy of Sciences - PNAS, 96(7), pp. 4084–4088.

Matei, V., Pauley, S., Kaing, S., Rowitch, D., Beisel, K. W., Morris, K., Feng, F., Jones, K., Lee, J. and Fritzsch, B. (2005) ‘Smaller inner ear sensory epithelia in Neurog 1 null mice are related to earlier hair cell cycle exit’, Developmental dynamics, 234(3), pp. 633–650.

Matsui, J. I., Gale, J. E. and Warchol, M. E. (2004) ‘Critical signaling events during the aminoglycoside-induced death of sensory hair cells in vitro’, Journal of neurobiology, 61(2), pp. 250–266.

McGovern, M. M., Hosamani, I. V., Niu, Y., Nguyen, K. Y., Zong, C. and Groves, A. K. (2024) ‘Expression of Atoh1, Gfi1, and Pou4f3 in the mature cochlea reprograms nonsensory cells into hair cells’, Proceedings of the National Academy of Sciences - PNAS, 121(5), pp. e2304680121–e2304680121.

Menendez, L., Trecek, T., Gopalakrishnan, S., Tao, L., Markowitz, A. L., Yu, H. V., Wang, X., Llamas, J., Huang, C., Lee, J., Kalluri, R., Ichida, J. and Segil, N. (2020) ‘Generation of inner ear hair cells by direct lineage conversion of primary somatic cells’, eLife, 9, pp. 1–33.

Mistry, B. A., D’Orsogna, M. R. and Chou, T. (2018) ‘The Effects of Statistical Multiplicity of Infection on Virus Quantification and Infectivity Assays’, Biophys J, 114(12), pp. 2974–2985.

Moore, S., Nakamura, T., Nie, J., Solivais, A., Aristizbal-Ramirez, I., Ueda, Y., Manikandan, M., Reddy, V. S., Romano, D., John, H., Perrin, B., Nelson, R., Frolenkov, G. and Hashino, E. (2023) ‘Generating high-fidelity cochlear organoids from human pluripotent stem cells’, Cell Stem Cell, 30(7), pp. 950–961.

Oliver, D., Knipper, M., Derst, C. and Fakler, B. (2003) ‘Resting Potential and Submembrane Calcium Concentration of Inner Hair Cells in the Isolated Mouse Cochlea Are Set by KCNQ-Type Potassium Channels’, The Journal of neuroscience, 23(6), pp. 2141–2149.

Phan, D. and Wodarz, D. (2015) ‘Modeling multiple infection of cells by viruses: Challenges and insights’, Math Biosci, 264, pp. 21–8.

Qiu, X., Mao, Q., Tang, Y., Wang, L., Chawla, R., Pliner, H. A. and Trapnell, C. (2017) ‘Reversed graph embedding resolves complex single-cell trajectories’, Nature methods, 14(10), pp. 979–982.

Ruben, R. J. and Sidman, R. L. (1967) ‘Serial Section Radioautography of the Inner Ear: Histological Technique’, Archives of Otolaryngology, 86(1), pp. 32–37.

Subramanian, A., Tamayo, P., Mootha, V. K., Mukherjee, S., Benjamin, L. E., Michael, A. G., Amanda, P., Pomeroy, S. L., Golub, T. R., Lander, E. S. and Mesirov, J. P. (2005) ‘Gene Set Enrichment Analysis: A Knowledge-Based Approach for Interpreting Genome-Wide Expression Profiles’, Proceedings of the National Academy of Sciences - PNAS, 102(43), pp. 15545–15550.

Tang, W., Ehrlich, I., Wolff, S. B. E., Michalski, A. M., Wölfl, S., Hasan, M. T., Lüthi, A. and Sprengel, R. (2009) ‘Faithful expression of multiple proteins via 2A-peptide self-processing: A versatile and reliable method for manipulating brain circuits’, The Journal of neuroscience, 29(27), pp. 8621–8629.

Vaisbuch, Y. and Santa Maria, P. L. (2018) ‘Age-Related Hearing Loss: Innovations in Hearing Augmentation’, Otolaryngologic clinics of North America, 51(4), pp. 705–723.

van der Valk, W. H., van Beelen, E. S. A., Steinhart, M. R., Nist-Lund, C., Osorio, D., de Groot, J. C. M. J., Sun, L., van Benthem, P. P. G., Koehler, K. R. and Locher, H. (2023) ‘A single-cell level comparison of human inner ear organoids with the human cochlea and vestibular organs’, Cell reports (Cambridge), 42(12), pp. 113527–113527.

Wiwatpanit, T., Lorenzen, S. M., Cantú, J. A., Foo, C. Z., Hogan, A. K., Márquez, F., Clancy, J. C., Schipma, M. J., Cheatham, M. A., Duggan, A. and García-Añoveros, J. (2018) ‘Trans- differentiation of outer hair cells into inner hair cells in the absence of INSM1’, Nature (London), 563(7733), pp. 691–695.

Wong, A. C. Y. and Ryan, A. F. (2015) ‘Mechanisms of sensorineural cell damage, death and survival in the cochlea’, Frontiers in aging neuroscience, 7, pp. 58–58.

Zinn, E. and Vandenberghe, L. H. (2014) ‘Adeno-associated virus: fit to serve’, Current opinion in virology, 8, pp. 90–97.

